# Biologically excretable AIE dots for visualizing through the marmosets intravitally: horizons in future clinical nanomedicine

**DOI:** 10.1101/2020.05.26.113316

**Authors:** Zhe Feng, Siyi Bai, Ji Qi, Chaowei Sun, Yuhuang Zhang, Xiaoming Yu, Huwei Ni, Di Wu, Xiaoxiao Fan, Dingwei Xue, Shunjie Liu, Ming Chen, Junyi Gong, Peifa Wei, Mubin He, Jacky W. Y. Lam, Xinjian Li, Ben Zhong Tang, Lixia Gao, Jun Qian

## Abstract

Superb reliability and biocompatibility equip aggregation-induced emission (AIE) dots with tremendous potential for fluorescence bioimaging. However, there is still a chronic lack of design instructions of excretable and bright AIE emitters. Here, we designed a kind of PEGylated AIE (OTPA-BBT) dots with strong absorption and extremely high NIR-II PLQY as 13.6%, and proposed the long-aliphatic-chain design blueprint contributing to their excretion from animal body. Assisted by the OTPA-BBT dots with bright fluorescence beyond 1100 nm and even 1500 nm (NIR-IIb), large-depth cerebral vasculature (beyond 600 μm) as well as real-time blood flowing were monitored through-thinned-skull, and noninvasive NIR-IIb imaging with rich high-spatial-frequency information gave a precise presentation of gastrointestinal tract in marmosets. Importantly, after intravenous or oral administration, the definite excretion of OTPA-BBT dots from the body was demonstrated, which showed an influential evidence of bio-safety.

## 1. Introduction

Exogenous photoluminescent organic dyes are widely perceived with good biocompatibility and have been extensively used in biopharmaceutics and imaging.^[1]^ However, luminophores always experience some effects of emission quenching, partially or completely in the state of aggregation, impeding the progress of some specific applications.^[2]^ In 2001, the uncommon luminogen system noted as aggregation-induced emission (AIE)^[3]^ broke down the captivity of Förster’s discovery named aggregation-caused quenching (ACQ), which brought a new wonderland for organic fluorophores. The intrinsic tendency to form aggregates in concentrated solutions or the solid state actively promotes the emission intensity of the fluorophores with AIE characteristics. Years of unremitting exploration has accumulated design experience of diverse AIEgens^[4]^ and shaped plentiful innovated applications for stimuli sensing,^[5]^ optoelectronic systems,^[6]^ molecular detection,^[7]^ bio-imaging^[8,9]^ and so on. Taking advantages of AIE dots with high resistance to photobleaching and excellent reliability, multifarious specific bio-sensing modes, including bio-imaging, launched on a grand ^[8,10]^

The development of AIE universe provides potential diagnostic and therapeutic means in clinic. Nowadays, efforts of AIEgens for fluorescence imaging in mice have already been made by many groups.^[8,11–14]^ As an essential link to advance clinically, the biosafety concerns of AIEgens were of vital importance but have not yet attracted sufficient attention. For instance, bio-excretion of AIE dots should doubtless be list in the guidance of next-generation AIEgens designs. Given the suppressed tissue-scattering and diminished autofluorescence, fluorescence bio-imaging in the second near-infrared region (NIR-II, 900–1700 nm) allows the precise visualization of details throughout some thick tissues in mammals with high signal-to-background ratio and sharp spatial resolution.^[15]^ It has been demonstrated that there existed a great improvement in imaging quality beyond 900 nm or even 1500 nm compared to the traditional NIR-I fluorescence imaging (760-900 nm),^[16]^ thereby gifting a credible approach for deep-penetration imaging *in vivo*. Hence, biological excretion, long emission wavelength and high emission intensity constitute the three harsh demands for new AIE dots, especially for imaging performed on primate models. Moreover, it should be noted that low doses of AIE dots might allow the rapid excretion from the organs, but also putting forward higher requirements for the dots’ emission intensity at the same time.

The common marmosets (Callithrix jacchus) are an influential and promising model for human behavior and diseases, and they were extensively applied in nanomedicine research due to its close physiologic, genetic and metabolic similarities to human beings.^[17]^ With its small body size and fast reproduction, limited amounts of test compounds are required to complete a program of studies and sufficient animals is provided to support research. The similarities between this species and humans in pharmacological activity, therapeutic targets, systemic exposure to drugs, kinetics, and metabolism have been proved by other scientific researchers.^[18]^ It is indispensable to understanding neural activity and cerebral blood flow regulation in marmosets for demystifying human brain diseases. In addition, the gastrointestinal (GI) structure of marmosets is analogue to that of human beings realistically,^[19]^ which greatly facilitates the researches about GI disorders. As an essential step towards clinic, the non-human primates study requires further exploration, and marmosets provide highly attractive models^[20]^ for the utilization of AIE dots in non-human primates imaging.

Herein, we discovered the long aliphatic chains of loaded AIEgens contributed significantly to the excretion of intravenously injected PEGylated micelles. Besides, we designed an excretable NIR-II AIEgen named as OTPA-BBT with a large molar absorption coefficient of ~5×10^4^ M^−1^ cm^−1^ at ~770 nm (in NIR-I spectral region). PEGylated OTPA-BBT dots with a peak emission wavelength of ~1020 nm and an extremely high NIR-II photoluminescence quantum yield (PLQY, measured as ~13.6%), were verified to emit ultra-bright luminescence beyond 1100 nm and even beyond 1500 nm (NIR-IIb). NIR-II fluorescence through-thinned-skull large-depth cerebrovascular microscopic imaging and NIR-IIb fluorescence noninvasive GI imaging in marmosets were well conducted for the first time by aid of the OTPA-BBT dots at incredibly low doses. Notably, the excretion of dots in hepatobiliary and gastro-intestinal pathway of non-human primates and rodents was substantiated drawing support from the sophisticated NIR-II fluorescence imaging technology. Thus, the potential application of the AIE dots on higher animals have been significantly pushed ahead and the eventual goal of clinical translation would be immensely accelerated.

## 2. Main text

### 2.1. Molecular design and photophysical properties

The fluorescence brightness of an emitter is determined by the absorption coefficient (◻) and PLQY (*Φ*), in which the former one reflects the ability of a chromophore to absorb light and the later expresses the efficiency to give out light via radiative decay. Therefore, it is of great interest to develop fluorophores with high ◻ and *Φ* simultaneously, yet these two parameters seem contrary with each other. From the molecular structure point of view, high ◻ usually requires a planar conformation to allow for efficient π-orbital overlap and oscillator strength, while high *Φ* is favorable in twisted structures to avoid the notorious ACQ effect. To solve this issue, a highly conjugated core structure with several twisting phenyl rings could be effective. The planar core unit with good conjugation would enable efficient electron transition, and the substituted benzene rotors are expected to inhibit intermolecular π-π stacking and afford AIE signature. Inspired by the previous work,^[21]^ we speculate that the aliphatic chains, influenced by some carrier protein, could help release the lipophilic molecules from micelles, which might contribute to the excretion of exogenous drugs *in vivo*. With this in mind, a strong donor-acceptor (D-A) type compound was built, in which octyloxy-substituted triphenyl amine (OTPA) and benzobisthiadiazole (BBT) served as the electron-donating and -withdrawing moieties, respectively, to design excretable, high brightness AIEgen with low bandgap (long-wavelength excitation/emission). The detailed syntheses and characterizations of OTPA-BBT and the intermediates are presented in Figure S1. Density functional theory (DFT) calculation suggests small dihedral angles (about 31°) between BBT and adjacent phenyl rings, indicating relatively planar molecular structure and good conjugation. While the peripheral phenyl derivatives are highly twisted, being beneficial for realizing AIE property. As displayed in **Figure** 1A, the electron density of the highest occupied molecular orbital (HOMO) is distributed on both the donor and acceptor moieties, whereas the lowest unoccupied molecular orbital (LUMO) is mainly localized on the electron-withdrawing BBT unit, indicating D-A characteristics with strong intramolecular charge transfer.

**Figure 1.**
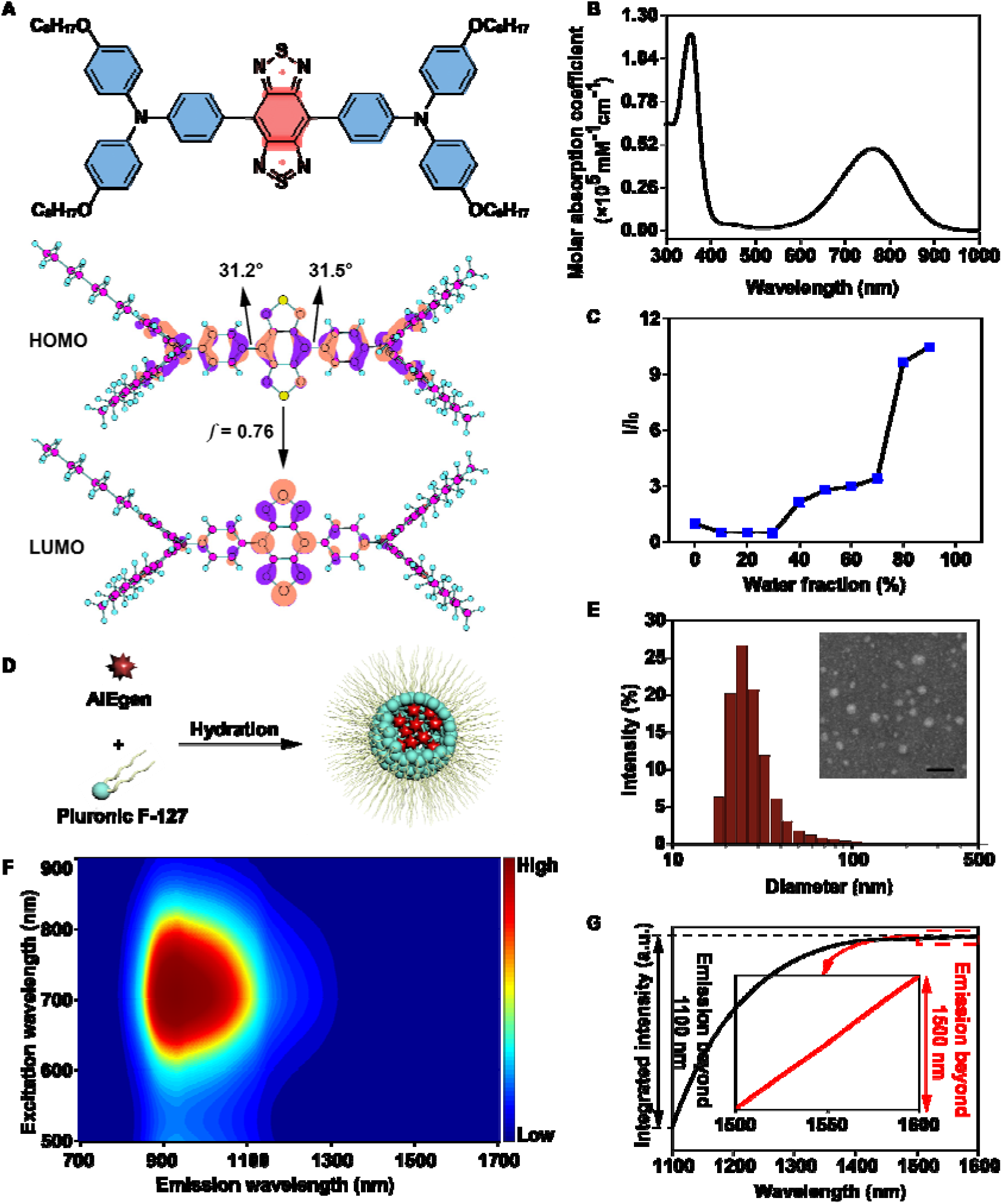
Photophysical properties of OTPA-BBT and the dots. A) The molecular structure (the red unit refers acceptor and the blue unit refers donator), as well as the calculated HOMO and LUMO wave functions. B) A spectrum showing the molar absorption coefficients of OTPA-BBT at various wavelengths. C) Plot of P.L. peak intensity of compound OTPA-BBT versus water fraction (*f*_*w*_) of THF/water mixture. I_0_ and I are the P.L. peak intensities of the OTPA-BBT in pure THF (*f*_*w*_ = 0) and THF/water mixtures with specific *f*_*w*_ s, respectively. D) Schematic illustration of the fabrication of OTPA-BBT dots. E) Representative DLS result and STEM image of the OTPA-BBT dots. Scale bar, 100 nm. F) P.L. excitation mapping of OTPA-BBT dots in aqueous dispersion. G) The integration of the fluorescence intensity. The value of the ordinate represents the integrated fluorescence intensity between 900 nm and the ordinate corresponding wavelength. The value difference on the ordinate represents the total fluorescence intensity in the corresponding spectral (abscissa) range. The insert image showed the integration of NIR-IIb emission.

The absorption spectrum of OTPA-BBT in THF is presented in Figure 1B. Interestingly, the compound exhibits broad absorption in NIR-I spectral region with a high molar absorption coefficient of 5 × 10^4^ M^−1^ cm^−1^ at ~770 nm, being advantageous for efficient light excitation. The oscillator strength (*f*) expresses the probability of a molecule to transit between different energy levels, representing an indicator of excitation ability. The *f* value of OTPA-BBT from HOMO to LUMO is calculated to be as high as 0.76, which suggests high electron transition possibility and explains the high ◻ value. We next investigated the photoluminescence (P.L.) properties in different aggregate states. P.L. intensity of OTPA-BBT gradually decreases by increasing the water fraction (*f*_*w*_) in THF/H_2_O mixture to 30%, which is probably due to the twisted intramolecular charge transfer (TICT) effect in high polarity environment. Then the emission intensifies when further increasing *f*_*w*_ to 90%, representing remarkable AIE signature (Figure 1C and S2A). The pronounced hypochromatic shift of maximal P.L. wavelength in high water fraction indicates the formation of aggregate and thus the polarity of microenvironment declines for the less possibility to contact with water molecule.

The AIEgens were then encapsulated into organic nanoparticles by nanoprecipitation,^[22]^ using US Food and Drug Administration (FDA)-approved F-127 (Figure 1D). The amphiphilic block copolymers could ameliorate the hydrophilicity and biocompatibility of luminescent probes. On the flip side, the matrix aggregated the AIEgens and the intramolecular motions was moderated by the pinning among the molecules. As shown in Figure 1E, the micellar systems were uniform with a narrow size distribution from dynamic light scattering (DLS, 28.6 ± 1.8 nm, Zetasizer Nano-ZS) and scanning transmission electron microscopy (STEM, 25.0 ± 0.8 nm, Nova nano 450). Taking IR-26 in dichloroethane as a reference (with a nominal PLQY = 0.5%), the NIR-II photoluminescence quantum yield of the OTPA-BBT dots was determined to be 13.6% (Figure S3). Due to the large absorption, high PLQY in NIR-II region and the nano-aggregation enhanced emission inside the micelles (Figure S4), the OTPA-BBT dots showed bright NIR-II fluorescence. The photoluminescence excitation (PLE) mapping was performed on the OTPA-BBT dots and demonstrated the excitation– emission relationships which brought us an in-depth understanding about the optical properties, as shown in Figure 1F. Furthermore, the integral spectrum of the fluorescence intensity in Figure 1G intuitively revealed the considerable emission and the discrepancy on the ordinate represented the total fluorescence intensity beyond 1100 nm and 1500 nm. Even though it took up only a tiny tail of the entire P.L. spectrum, especially beyond 1500 nm, the signals could still be well sensed by the InGaAs camera (Figure S5), which foreshadowed the great potential for NIR-IIb (beyond 1500 nm) fluorescence imaging. The OTPA-BBT dots suspended in phosphate buffer saline (PBS), serum, simulated intestinal fluid and gastric fluid were then exposed to 793 nm continuous-wave (CW) laser irradiation at a power density of 120 mW cm^−2^ for 30 min, whose power was higher than that during realistic imaging *in vivo*. As shown in Figure S6, the superior photostability and chemical stability in the extreme environments could be recognized. In addition, there is no perceptible difference in morphology, particle size and optical properties of dots after 5-day dispersion in acidic (pH = 2) and alkaline (pH = 12) media (Figure S7). The OTPA-BBT dots could also be successfully freeze-dried and rehydrated, as shown in the Figure S8, illustrating the medicament has the potential of long-term storage and even industrialization development in the future.

### 2.2 *In vivo* bio-excretion and bio-distribution of the AIE dots

The toxicity and excretion of exogenous nanomaterials introduced into a body are of important concerns. Over a period of 4 h post intravenous injection (2 mg kgBW^−1^) of OTPA-BBT dots, the signals in liver and intestines evidently accumulated, demonstrating the uptake from blood circulation (**Figure** 2A and S9A). We collected the feces excreted from mice, recorded the fluorescence and observed the strong emission of the OTPA-BBT dots. As shown in Figure 2A and S3A, as expected, the excretion by the hepatobiliary pathway resulted in a dramatical intensity decrease of the signals in the organs within 44 days post injection, especially liver and intestines. The fluorescence intensity of the isolated liver and spleen of mice 6 weeks post injection was also measured, which showed remarkable decrease (at least 98% and 97%, respectively) compared to that at 1st week post-injection (Figure S9B). On the 1st day and 28th day post injection, the biochemical test and blood routine test results showed that the OTPA-BBT dots did not influence the hepatic and renal function, or peripheral blood cells (Tables S1-S8), and the results of histology study of major organs showed no obvious side effects of OTPA-BBT dots *in vivo* (Figure S9C). So far, no evidence showed appreciable toxicity of the OTPA-BBT dots *in vivo*, which could facilitate the pre-clinical trials in non-human primate.

**Figure 2.**
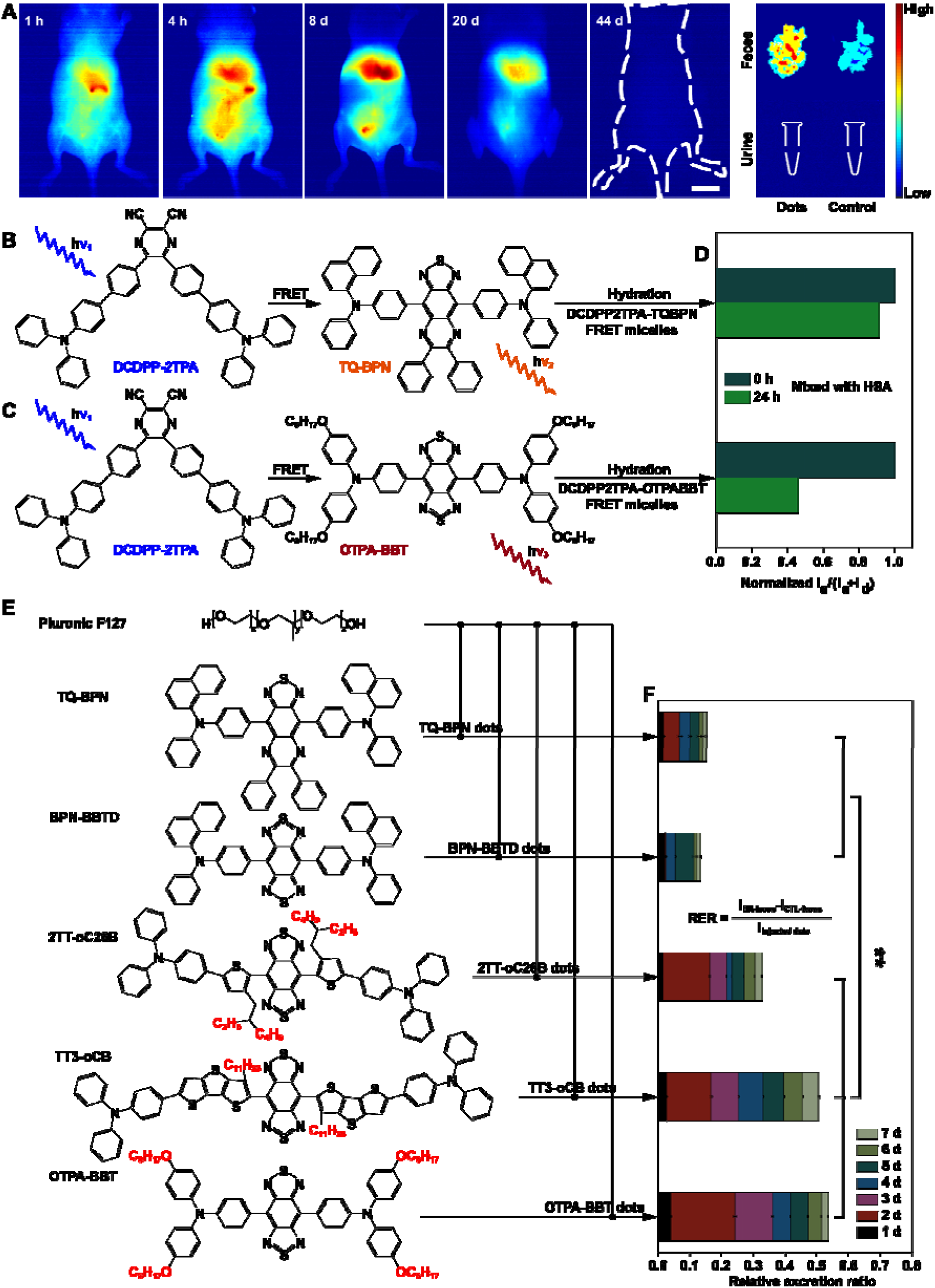
*In vivo* bio-excretion of several kinds of AIE dots. A) The excretion monitoring of intravenously injected OTPA-BBT dots in mice (2 mg kgBW^−1^), via NIR-II fluorescence imaging. Scale bar, 10 mm. B) The energy overlap between DCDPP-2TPA (donor) and TQ-BPN (acceptor). C) The energy overlap between DCDPP-2TPA (donor) and OTPA-BBT (acceptor). D) The traces of normalized FRET ratio at 0 h and 24 h post-mix. E) Schematic illustration of several kinds of AIEgens loaded PEGylated micelles. F) The relative excretion ratio of the five kinds of AIE dots in 7 days post intravenous injection (10 mg kgBW^−1^). ** indicates p < 0.01 for T-test.

Metabolism of exogenous drugs *in vivo* is a complex biological process, in which some proteins are always associated with docking, transport, and excretion (e.g., the excretion of aspirin). Previous work has indicated that PEGylated micelles could release the lipophilic agents^[21]^ and we speculated the long carbon chains of the loaded agents might play a certain role. For further analysis, fluorescence resonance energy transfer (FRET) technology was utilized to monitor the release of micelles. A kind of AIEgen, named DCDPP-2TPA,^[23]^ was chosen as donor. Meanwhile, TQ-BPN, a kind of reported AIEgen without aliphatic chain,^[14]^ and OTPA-BBT with four aliphatic chains were selected as acceptors respectively (Figure S10A and S10B, Figure 2A-2C). After hydration, two lipophilic FRET pairs (DCDPP2TPA-TQBPN and DCDPP2TPA-OTPABBT) were physically entrapped into micelle cores. The two kinds of FRET micelles were then respectively mixed with human serum albumin (HSA), a widespread protein in the blood as well as liver that could transport fatty acids, and the emission spectra and FRET ratios I_a_/(I_a_ + I_d_) (Subscript ‘a’ refers to acceptor and ‘d’ refers to donor) were measured after 0 and 24 hours. As shown in Figure S10C, S10D and Figure 2D, in contrast of DCDPP2TPA-TQBPN dots (the core-loaded molecules, DCDPP-2TPA and TQ-BPN, had no long aliphatic chains), the obvious reduction in FRET efficiency of DCDPP2TPA-OTPABBT dots confirmed the release of core-loaded molecules (OTPA-BBT) with long aliphatic chains. The disintegration of AIE dots should be the first step, and then proteins that affected the release of AIE molecules from micelles could further stimulate their excretion.

To detect the excretion of AIEgens with aliphatic chains *in vivo* directly, we selected four other typical kinds of NIR-II AIE fluorophores in which the two possess aliphatic chains (2TT-oC26B^[13]^ and TT3-oCB^[24]^) but the others (TQ-BPN^[14]^ and BPN-BBTD^[11]^) do not. With OTPA-BBT (in this work), we PEGylated all the five kinds of AIEgens via Pluronic F127 (Figure 2E) and intravenously injected them to the mice respectively (10 mg kgBW^−1^). The feces were collected daily for 7 days and the relative excretion ratios were calculated by the fluorescence intensities of the injected dots (I_injected dots_) and the feces in the experiment groups (I_EX-feces_) as well as the control groups (I_CTL-feces_). According to Figure 2F and Tables S9-S10, significant differences (T-test, p = 0.002) between the dots loading the AIEgens with (2TT-oC26B, TT3-oCB, OTPA-BBT dots) and without (TQ-BPN, BPN-BBTD dots) aliphatic chains were indicated. As shown in the DLS results in Figure S11, the particle sizes of the dots were about dozens of nanometers and STEM images illustrated the five kinds of dots possessed similar spherical shapes. It could be concluded that the long aliphatic chains could be conducive to the release of AIE molecules from PEGylated micelles and have a marked impact on the further excretion of them from body, which ought to be a valuable element for guiding the design of novel excretable molecular agent in future.

### 2.3 High-spatial-resolution through-thinned-skull cerebrovascular microscopic imaging

In the fields of biological research, the deciphering of cerebrovasculature was inextricably linked to the exploration of neuroinformatics and brain-related theory in rodents, humans and other species.^[25]^ Meanwhile, some common neurological disorders such as Alzheimer’s or amyotrophic lateral sclerosis may be related to dysfunction of cerebrovascular structure and/or function.^[26]^ In our study, OTPA-BBT dots at a low dose (2 mg kgBW^−1^) could render the cerebral vascular network under the cranial window of mice clearly visible (Figure S12A). By axially tuning the relative position between the sample stage and the objective lens, the signals on each focal plane within nearly 870 μm below thinned-skull were almost thoroughly recorded and an ultra-high spatial resolution of ~2.4 μm was attained uncomplicatedly (Figure S12B- S12M).

The observation of brain vasculatures was then extended to marmosets and we expanded on the feasibility of harmless OTPA-BBT dots assisted NIR-II fluorescence imaging of the cerebrovascular structure in marmosets, New World monkeys (**Figure** 3A). NIR-II fluorescence wide-field microscopy is characterized as easy operation and minute loss of effective signals. In order to gather more NIR-II signals through the thinned skull, we collected fluorescence emission beyond 1100 nm in vessels. After intravenous injection of the OTPA-BBT dots (2 mg kgBW^−1^), a large view of cortical vasculature was preserved on the NIR-sensitive camera by a scan lens (LSM03, Thorlabs). Tracking the periodic vibrations of the capillaries, we measured the respiratory rate (0.64 Hz) and heart rate (3.30 Hz) of the marmoset in time (Figure 3B-3E and Movie S1, the other two peaks at 1.28 Hz and 6.60 Hz in Figure 3D and 3E represent the second harmonics of heart rate and respiratory rate respectively). The calculation is within normal range and well-echoed the actual values measured by an ECG (electrocardiogram) monitor (Figure S13A). We also focused on the excretion of OTPA-BBT dots from marmosets and the P.L. intensities of metabolites (feces and urine) collected at certain time points were verified post-injection. As shown in Figure 3F and Tables S11-S16, the hepatobiliary excretion route in marmosets was confirmed and no noticeable effects of the OTPA-BBT dots on the hepatic and renal function, or peripheral blood cells have been tested, which were consistent with that in mice. Next, a typical micro cortical region was locked and the NIR-II through-thinned-skull tomographic angiography in brain was further conducted with a 25X objective. Interestingly, imaging depth below thinned-skull reached nearly 700 μm eventually and a capillary of ~5.2 μm (diameter) at ~200 μm was distinctly identified, which bettered previous records, owing to the extraordinarily bright OTPA-BBT dots (Figure 3G-3N).

**Figure 3.**
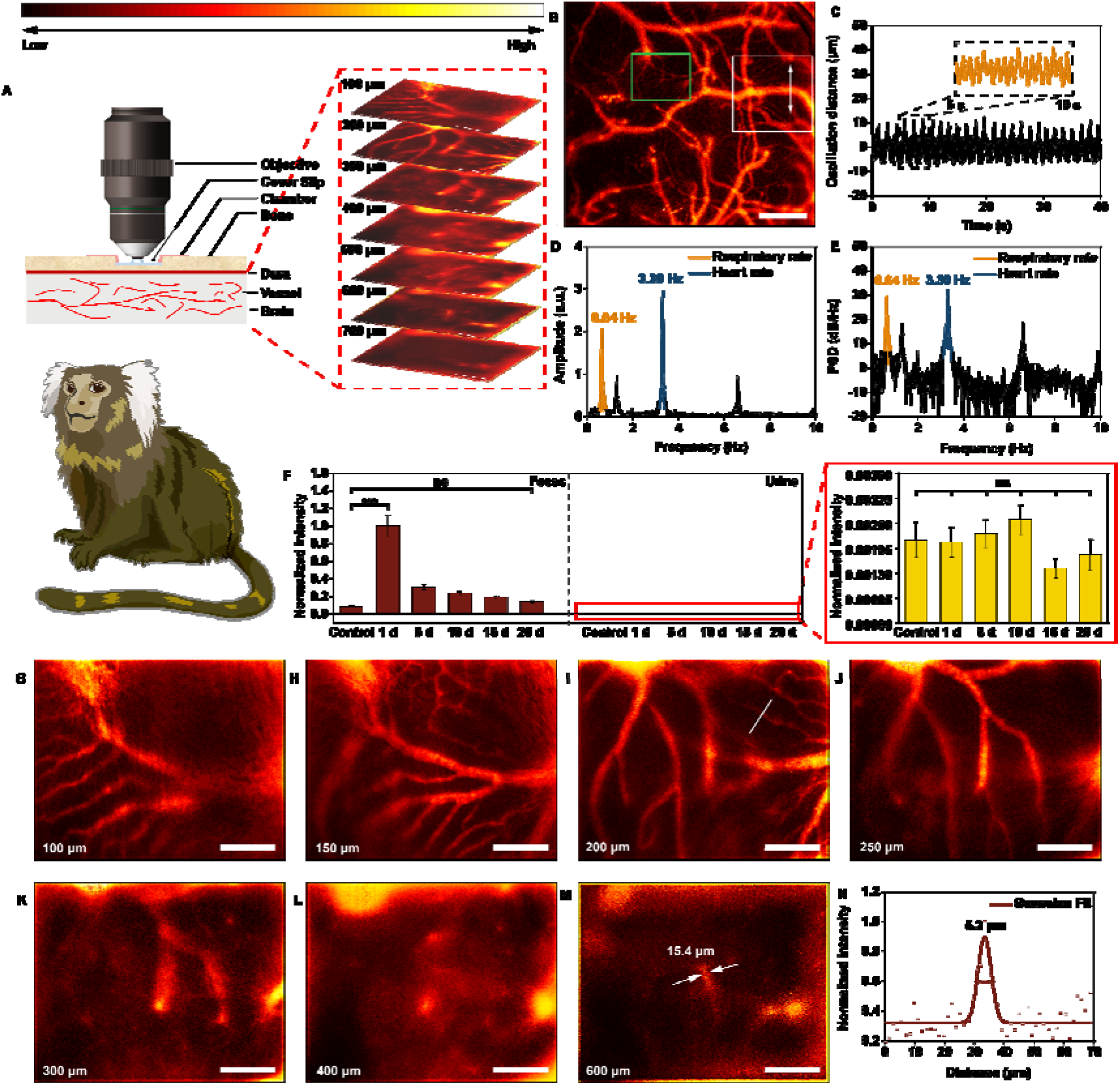
Through-thinned-skull cerebrovascular microscopic imaging with high spatial-resolution. A) The operation illustration of microscopic imaging. The skull of the marmoset was just thinned rather than opened, aiming at minimal injury for bone regeneration. B) A NIR-II fluorescence microscopic image (~5X) of brain vasculature. Scale bar, 300 μm. C) The position of a cortical vessel was plotted as a function of time. D) The fast Fourier transformation of time-domain signals in (C). E) The power spectral density of (C). F) The normalized P.L. intensity of the feces and urine from the marmosets post intravenous injection of OTPA-BBT dots (2 mg kgBW^−1^). Error bars indicate s.e.m. (n = 5, T-test, *** P< 0.001, ns P > 0.1). G) - M) Images at various depths below the thinned skull. Scale bar, 100 μm. N) Cross-sectional fluorescence intensity profile along the white line of the cerebral blood vessel (inset of (I)). The Gaussian fit to the profile is shown in red line in (N), which showed a high spatial resolution.

### 2.4 High-temporal-resolution through-thinned-skull cortical blood flow monitoring

The flow and circulation of blood maintain the normal operation of body. An efficient method for real-time cerebrovascular monitoring means a great deal in the study of life sciences. In a fixed field of view, cortical vasculatures in marmosets with diverse functions always possess different rates. Assisted by the bright emission and biocompatibility of OTPA-BBT dots, the adrift fluorescent spots in blood could be readily captured by the microscope through-thinned-skull. We recorded the regions of interest continuously per 40 ms and the positions of the points as a function of time were plotted (**Figure** 4A-4B and Figure S13B-S13G). In each blood capillary, the calculation was repeated three times to obtain the mean flow rate of the six blood vessels in the selected area (Figure 4C). Additionally, the type of blood vessels can be classified as veins based on the direction of flow, as veins collect blood from the branches, whereas arteries do the opposite (Movie S2).

**Figure 4.**
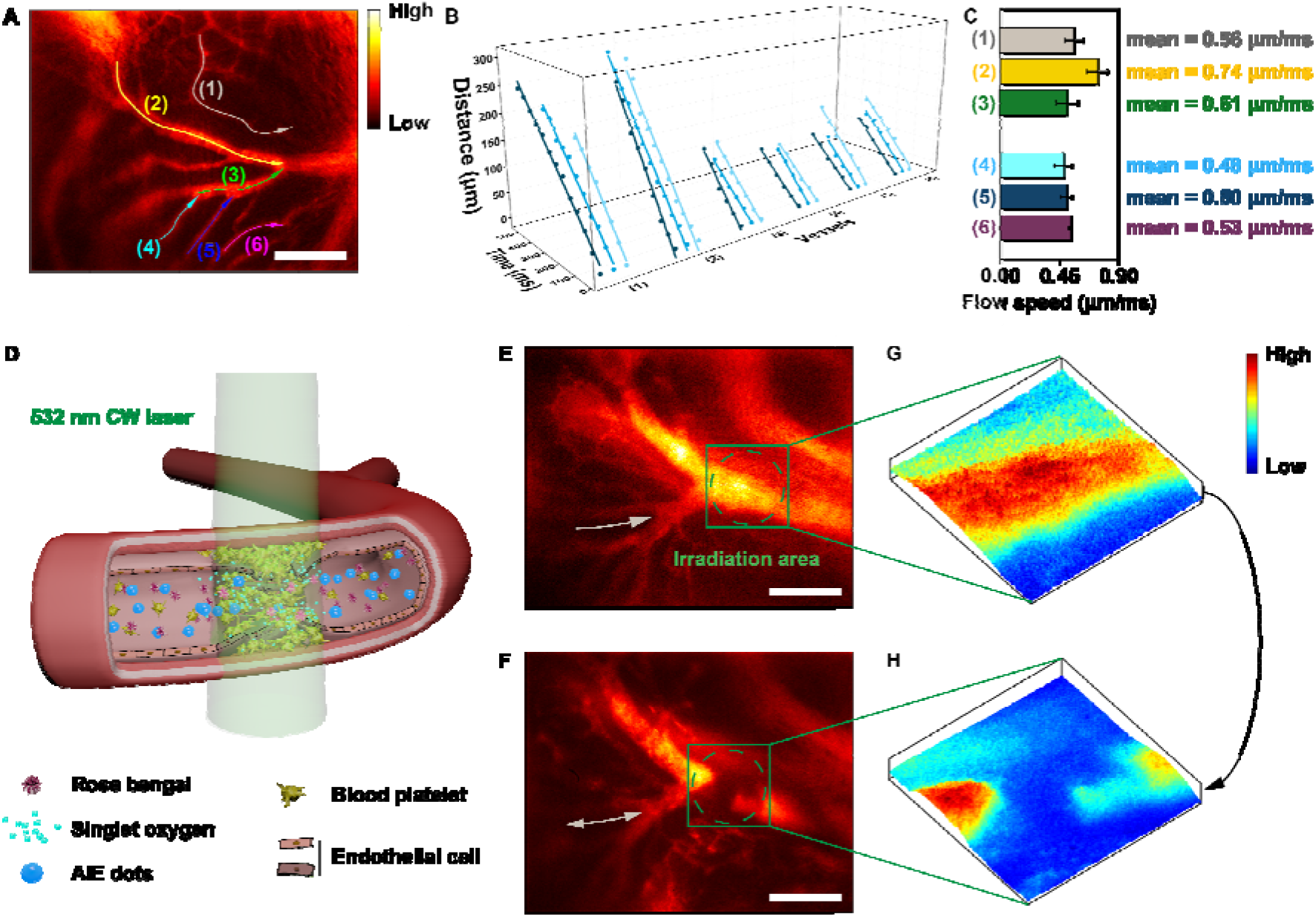
Through-thinned-skull cortical blood flow monitoring with high temporal-resolution. A) A microscopic image (~25X) of brain vasculature and six vessels were selected. Scale bar, 100 μm. B) The plots of positions of the fluorescent point signal as a function of time (3 times) in the vessel (1) - (6) in (A). C) The mean blood flow rates in the vessel (1) - (6) in (A). Error bar, s.e.m. across trials. D) Schematic illustration of PTI induction under excitation of 532 nm CW laser. E) - F) NIR-II fluorescence microscopic images of brain blood vessels before (E) and after (F) PTI induction. Scale bar, 100 μm. G) - H) 3D NIR-II fluorescence intensity distributions of the irradiation area before (G) and after (H) PTI induction. The white arrows stand for the directions of flow before and after PTI induction. PTI induced the alteration of the blood flow direction.

The aftermaths of a stroke primarily depend on the location of the obstruction and the extent of related brain tissue, and it could lead to the alteration of blood flow direction. NIR-II fluorescence wide-field microscopic imaging could provide a noninvasive method for thrombosis monitoring (Figure 4D). The photothrombotic induction of capillary ischemia was observed firstly in mice with cranial windows. Figure S14A showed a cerebrovascular image with large field-of-view post injection of OTPA-BBT dots, in which a region of ~454 μm × ~363 μm was selected (Figure S14B). After the Rose Bengal was injected into the mouse, the area outlined by the green dashed circle was continuously illuminated for 150 seconds by 532 nm laser post injection of photosensitizer. Figure S14C revealed that the stockpiled platelets impeded the blood flow, making the dots unable to circulate continuously. Before long the slight obstruction was flushed away by the high-speed blood flow that contained OTPA-BBT dots and the two once-occluded capillaries returned to normal in ~230 s (Figure 14C- S14E, Movie S3 and Movie S4). Next, we shrank the irradiation area to increase energy density in another mouse. After irradiation for 150 s, there was no obvious recovery of blood flow in the next 60 min, while some vascular hyperplasia could be nevertheless observed (Figure S14F-S14K). Moreover, the photothrombotic thrombosis showed multiple effects on nearby blood vessels, and the vascular morphology changes were shown in Figure S14L and S14M.

Real-time through-thinned-skull functional imaging in brain assisted by the OTPA-BBT dots was further conducted on marmosets, and cerebrovascular alteration after brain embolism was observed on marmosets, giving another noninvasive diagnostics modality for primates. Thanks mainly to the ultrahigh-brightness of the OTPA-BBT dots, the blockage of cortical vasculatures below the thinned skull was exceedingly visible post irradiation of the 532 nm CW laser for 3 min (Figure 4E-4H). Interestingly, the real-time video of the ROI wrote down the blood flow arrest or even reflux in other side branches (Movie S5).

### 2.5 NIR-IIb fluorescence high-spatial-frequency noninvasive gastrointestinal imaging

Pathologies of GI tract such as neoplastic and functional pathologies, chronic inflammatory diseases, and nontraumatic emergency causing occlusion are multifarious and patients of diverse ages can be affected by such diseases. Consequently, clinically common diagnostic approaches, like computerized tomography (CT) or magnetic resonance imaging (MRI), should be reassessed seriously in view of the ionizing radiation, limited spatial resolution or time consuming.^[27]^ NIR-IIb (1500-1700 nm) fluorescence imaging with the extremely low scattering overcomes the above shortcomings to a certain degree and stands out in optical biological imaging on account of its superlative penetration depth in complex refractive index media. Thus, it is of great clinical value in functional diagnosis of the GI tract.

As a suitable contrast agent, the OTPA-BBT dots in our work was demonstrated possessing excellent stability and strong NIR-IIb emission. After intragastric administration (15 mg kgBW^−1^), the NIR-IIb fluorescence GI imaging was carried out in mice. During the first 20 minutes, the fluorescent probe passed into the jejunum from the duodenum and the ileum was lighted up progressively within 1 h post-treatment (Figure S15A). The excretion from ileum to colon was also recorded throughout the next 9 h (Movie S6). The typical image at 22 h precisely showed the feces that have not yet been excreted thoroughly. In contrast to the increasingly blurry details imaged in the NIR-II region beyond 1300 nm, 1200 nm, 1100 nm, and 1000 nm (Figure S15B and S15C), the sharpest resolution and highest signal-to-background ratio (the background was defined as the valley between the two peaks) images of ileum were obtained beyond 1500 nm, and we could see that much sharper images were got by extending the wavelength of the detected photons. After 32 h, the feces almost gathered all the fluorescent emission. Importantly, the main organs remained no fluorescent signals at all at 3 d post-treatment and the safety of the OTPA-BBT dots was demonstrated (Figure S15D-S15F).

Next, we evaluated NIR-IIb GI imaging in marmosets, which might contribute on the clinical applications in the future. The operation illustration was shown in **Figure** 5A and the sharp NIR-IIb images of marmosets deciphered the complicated bowel structure well (Figure 5B-5M, Figure S16 and Movie S7) post feeding (15 mg kgBW^−1^). At 60 min, we analyzed the small intestine structure in the traditional NIR-II (beyond 1000 nm), the NIR-IIa (beyond 1300 nm), and the NIR-IIb (beyond 1500 nm). As shown in Figure 5G, the gap between intestines was accurately identified as 1.04 mm without being submerged in fluorescence, benefiting from the sharp edges in the NIR-IIb image and the stable bright emission of the probe. To further figure out the superiority of NIR-IIb fluorescence imaging, we singled out the representative parts separately from the images beyond 1000 nm, 1300 nm, and 1500 nm and their spectrum maps were shown in Figure 5H-5J by fast Fourier transformation (FFT) (Figure S17). The fact that the NIR-IIb image is in possession of the most abundant high-frequency information was confirmed, that is to say, the imaging technology beyond 1500 nm possesses precise expression of details and edges. As expected, the OTPA-BBT dots were also excreted smoothly from the marmosets eventually (Figure 5N and Tables S17-S18). This work is the first attempt to apply NIR-IIb fluorescence imaging technology in primate mammals, and we believe it holds immeasurable significance.

**Figure 5.**
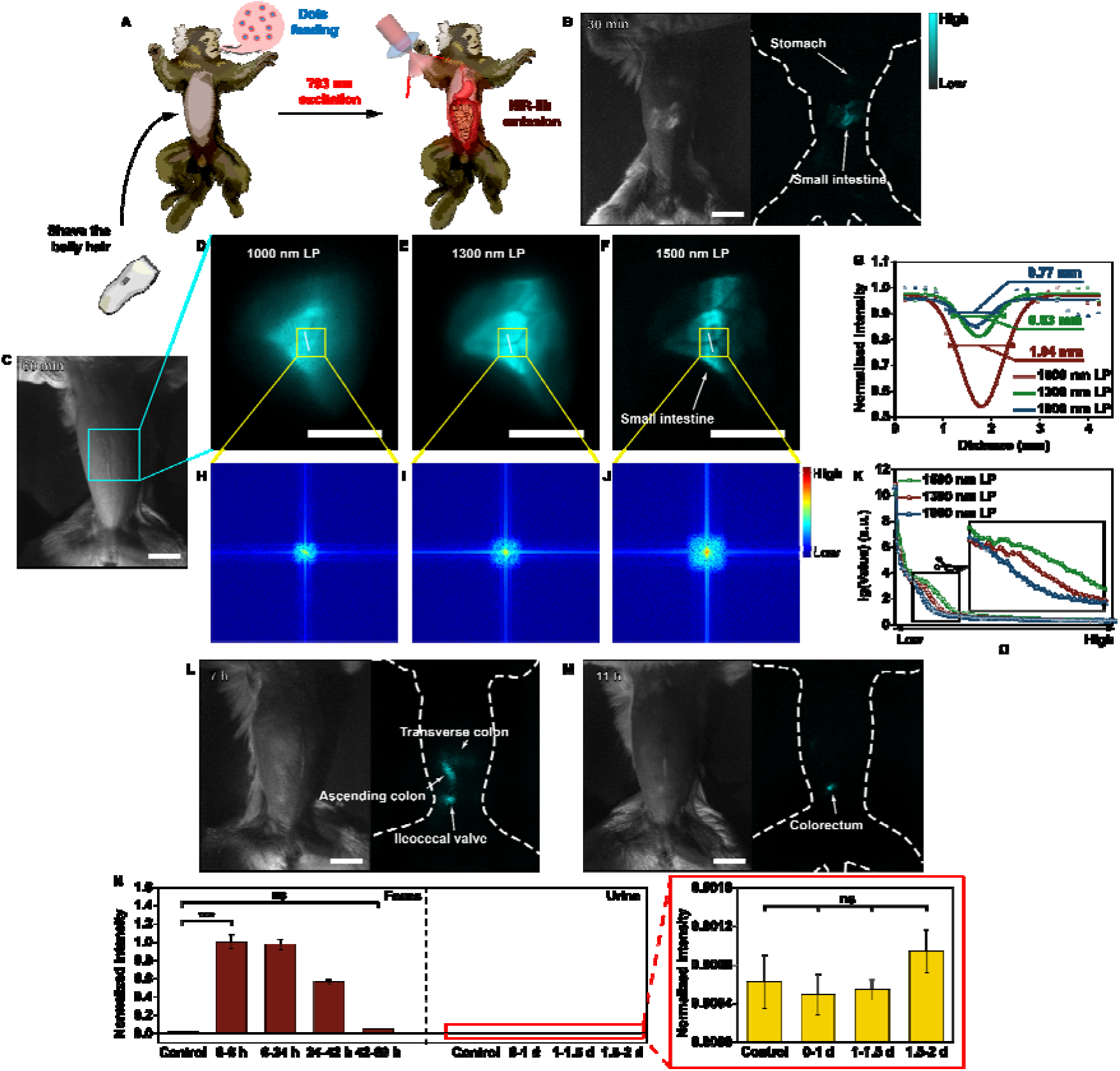
NIR-IIb fluorescence noninvasive gastrointestinal imaging with high spatial-frequency. A) The operation illustration of GI imaging. B) The NIR-IIb fluorescence GI images at 30 min post feeding. C) - F) The bright-field, NIR-II (beyond 1000 nm), NIR-IIa (beyond 1300 nm), and NIR-IIb (beyond 1500 nm) fluorescence GI images at 60 min. G) Cross-sectional fluorescence intensity profiles along the white line of the intestines (inset of (D), (E) and (F)). The Gaussian fit to the profile is shown in blue (D), green (E) and red (F) line in (G). The NIR-IIb image showed better resolution capability. H) - J) The FFT images of (D), (E) and (F). K) The frequency distribution of (H), (I) and (J). The NIR-IIb image was confirmed to hold more high-frequency information. Scale bar, 20 mm. L) and M) The NIR-IIb fluorescence GI images at 7h and 11h post treatment. N) The normalized P.L. intensity changes of the feces and urine from the marmoset post feeding of OTPA-BBT dots (15 mg kgBW^−1^) over time (mean ± s.e.m., n = 3 independent samples, T-test, *** P < 0.001, ns P > 0.1). Scale bar, 20 mm.

## 3. Conclusion

Aggregation-induced emission has been respected as a major research focus, since the uncommon emission behaviors were revealed around two decades ago. Low toxicity and good biocompatibility equipped these organic fluorophores with huge potential for biological applications. Biological excretion and bright emission are the given direction for novel NIR-II fluorescent AIEgens design. We found and proved that the long carbon chains of the loaded AIEgen significantly enhanced the excretion of the dots from body. In view of this, we developed PEGylated OTPA-BBT dots exhibiting typical AIE characteristics. The dots were proved undetectably bio-toxic and well biocompatible. The strong emission of marmoset dejecta testified the obvious hepatobiliary excretion within 26 days post intravenous injection. Relying on the ultrabright OTPA-BBT dots, noninvasive through-thinned-skull cerebrovascular microscopic observation was conducted with high spatial and temporal resolution in marmosets. It is encouraging that the OTPA-BBT dots we reported exhibited considerable fluorescence even beyond 1500◻nm, bringing us a versatile NIR-IIb dye capable of ultrahigh-definition GI imaging in non-human primates.

Marmosets are acknowledged as subjects in studies involving toxicology, reproductive biology, neurology, behavior, cognition, and more, thus always taking huge responsibility for human medical research. We took the lead to apply biocompatible OTPA-BBT dots to marmoset fluorescence imaging, and splendidly performed through-thinned-skull cortical vasculature imaging and NIR-IIb fluorescence GI imaging in non-human primates for the first time. Future research will focus on the exhaustive toxicity investigation of the OTPA-BBT dots on primates. We think that the work could to some extent guide next-generation AIEgens designs and promote the development of AIEgens in large animal application.

## 4. Experimental/Methods Section

### 4.1 PEGylation of AIEgens

1 mL of tetrahydrofuran (THF) solution containing 2 mg of OTPA-BBT/TQ-BPN/2TT-oC26B/TT3-oCB AIEgen and 10 mg of Pluronic F-127 was poured into 10 mL of deionized water, followed by sonication with a microtip probe sonicator (XL2000, Misonix Incorporated, NY) for 5 min. The residue THF solvent was evaporated by violent stirring the suspension in fume hood overnight, and colloidal solution was obtained. 10 mg of Pluronic F-127 was mixed with 2 mg BPN-BBTD AIEgen in 2 mL of chloroform and sonicated for 5 min. The mixture was then dried under vacuum in a rotary evaporator at 40 °C. After the chloroform was evaporated completely, 10 mL of deionized water was added to dissolve the residue. The micelles (OTPA-BBT/TQ-BPN/2TT-oC26B/TT3-oCB/BPN-BBTD dots) were then subjected to ultrafiltration for further purification.

As for the FRET dots, the mass ratio of donor to acceptor is set to be 1:1. 1 mL THF solution of containing 1 mg of TQ-BPN/OTPA-BBT AIEgen, 1 mg of DCDPP-2TPA AIEgen and 10 mg of Pluronic F-127 was poured into 10 mL of deionized water, followed by sonication for 5 min. The colloidal solution was obtained by violent stirring the suspension in fume hood overnight and evaporating the residue THF solvent. The FRET dots (DCDPP2TPA-TQBPN and DCDPP2TPA-OTPABBT dots) were then subjected to ultrafiltration for further purification.

### 4.2 Animal handing

All experimental procedures were approved by Animal Use and Care Committee at Zhejiang University and in accordance with the National Institutes of Health Guidelines. Experimental animals: Mice (BALB/c and nude mice) and common marmosets (Callithrix jacchus). We used young marmosets (150-200 g), 6~7 months of age, with either sex in this study. BALB/c and nude mice were fed with water and food with a normal 12 h light/dark cycle, and the surrounding relative humidity level was 55–65% and the temperature was ~25 °C. Marmosets were generally raised in a suitable environment of about 26 °C, the indoor temperature was kept constant, and the relative humidity was around 40%~70%. All groups within study contained n = 10 mice and n = 4 monkeys.

### 4.3 Surgery for cerebrovascular microscopic imaging of marmosets

Marmosets (Callithrix jacchus, ~200 g) were initially anesthetized with ketamine (30 mg kgBW^−1^) by intramuscular injection and maintained with 0.5–3.0% isoflurane in 100% medical oxygen by a facemask connect to an inhalation anesthesia machine (RWD 510). The animals were placed in a marmoset stereotaxic frame (SR-5C-HT, Narishige) and the body temperature was maintained by a thermostatic heating pad at 37°C. The heart rates (180–300 bpm) and respiration rates (25–40 breaths per minute) were monitored and arterial oxygen saturation (SpO_2_) was kept above 95% (v/v) by adjusting the concentration of isoflurane. The head of the animal was shaved with hair clippers and scrubbed with three washes of 10% (v/v) povidone iodine, followed by 70% (v/v) ethanol. All additional surgical steps were conducted under aseptic or sterile conditions. A midline incision was made to expose the skull. A craniotomy with 5.0 mm in diameter was grinded to ~50% of the thickness of the bone in a target imaging site using a mini handheld cranial drill (RWD 78001). A round glass coverslip was placed on the top of the craniotomy and secured by cyanoacrylate adhesive and dental cement, respectively, to create an observing window for the cerebral vessels imaging later. A wall was built surrounded the glass coverslip with dental cement to hold deuterium oxide linking the cranial window and objective. The unnecessary glue covering the coverslip was carefully removed by using blunt side of scalpel under an operating microscope. After the imaging sessions were finished, the coverslip was removed from the skull and the wounds were flushed with fresh saline. Finally, sterilized sutures were tied to repairing a large surgical defect on the scalp along the midline of the skull. Animals were administrated with ceftriaxone (22 mg kgBW^−1^) for the subsequent 7 days to minimize inflammation and infection.

### 4.4 Procedures for cerebrovascular microscopic imaging of marmosets

After anaesthesia, the marmosets were immobilized on a two-dimensional (2D) translational stage with adjustable X and Y axis, placing the region of interest right under the objective lens (Figure S18A). Then 10 cc warm glucose in normal saline and OTPA-BBT dots (2 mg kgBW^−1^) were administrated by intravenous catheter, respectively. As shown in Figure S18B, a collimated 793 nm CW laser was introduced into the modified epi-illumination (NIRII-MS, Sunny Optical Technology, China), expanded by lens, reflexed by a long-pass dichroic mirror and focused on the back focal plane of the objective lens, eventually irradiating the sample evenly and securely. The excitation light from the objective lens penetrated the thinned skull and dura to light up the dots in the blood vessels. In turn, the emission signals beyond 1100 nm were collected by the objective lens through the thinned skull and the dichroic mirror, and afterwards focused on an NIR-sensitive camera (SW640, Tekwin, China). To purify the NIR-II signals, the long-pass optical filter was set in front of the 2D-detector. During imaging, the images at increasing depths were recorded by vertically tuning the epi-illumination.

### 4.5 Procedures for photothrombotic induction

As shown in Figure S18C, a 532 nm CW laser beam was selected as the excitation source and introduced into the wide-field microscope for the photothrombotic induction. After collimated, reflected and focused, it eventually irradiated evenly the sample with tiny area for a local photodynamic damage (~20 mW before the objective). After intravenously injected (20 mg kgBW^−1^), the photosensitizer (Rose Bengal) could release singlet oxygen (^1^O_2_) under excitation that then adhered to and destroyed the endothelial cells of blood vessels. Platelets those spontaneously stuck together in damaged regions could then form clots, which would eventually yield cerebral strokes. Thrombus was induced and monitored by switching the excitation source and equipped optical set-up.

### 4.6 Procedures for gastrointestinal imaging of marmosets

Marmosets were trained to be fed by syringe without wasting food for 3-5 days before GI imaging experiment. During the imaging sessions, animal was fed with OTPA-BBT dots (15 mg kgBW^−1^) by syringe. The animal was anesthetized with ketamine (30 mg kgBW^−1^) by intramuscular injection and placed on a height-adjustable table with belly up after the hair on the bell was shaved. As shown in Figure S18D, the expanded laser (793 nm, 120 mW cm^−2^) illuminated the whole belly and a fixed focus lens (TKL35, Tekwin, China) collected the emission signals from GI tract. The belly of the animal was imaged for 30 s every 5 minutes in the first thirty minutes and every 2 hours in the following time of the experiments. While the animal was in awake state, we hold the animal still with belly up for 30 s for imaging by leather gloves. The animal was allowed returning to the home cage for eating and drinking in the imaging interval of the experiments. After GI imaging experiment was finished, the experimental animals were raised in single cage for 3-5 days, which is convenient for collecting the urine and feces.

### 4.7 Image signal was processed by 2-D Fourier transform

Low-frequency components in the spectrum correspond to areas with flat gray scale variations in the image space, where energy is mainly concentrated. Nevertheless high-frequency components correspond to areas with sharp grayscale variations, which are principally represented by edges and details in the original image.

After an image undergoes Discrete Fourier transform (DFT), the low-frequency components are reflected in the four corner parts of spectrum. To facilitate the observation of the spectrum distribution, the low-frequency components are shifted to the spectrum center through the spectrum centralization (Figure S17) and the outer part of spectrum is basically the high frequency component. As shown in Figure 5D-5F, the more high-frequency information an image contains, the more detail and clarity it contains.

### 4.8 Excreta collection and postoperative observation

We administrated the AIE dots onto animals’ bodies via intravenous catheter for the experimental group and sterile distilled water for control group. Both groups were raised in the same environments. We collected the metabolites from both groups daily or one certain day per week for 6-8 weeks. In the GI imaging experiment, we collected feces and urine right after the animals was fed with fluorescent dots. The collection was lasted for 3-5 days in the experimental animals, while collecting excretion from the animals in the control group fed with sterile distilled water, and there should be no contact between urine and feces during collection. The marmosets’ body temperature, body weight, food intake, and activity levels were recorded before and for two months after the experiment. The recorded results showed that the body weight of three marmosets varied within a normal range. Additionally, there was no obvious abnormality in activity levels and basic exploratory behavior of experiment marmosets.

### 4.9 Statistical Analysis

All data analyses were shown as mean ± standard errors of the mean. The data of fluorescence intensity between two groups with normally distributed and equal variances were analyzed by calculating two-tailed independent sample T test. P values < 0.05 were considered statistically significant for all the analyses and were indicated with asterisks, such as * P < 0.05, ** P < 0.01, *** P < 0.001. Detailed information including sample size and statistical differences between groups was shown in Tables S10 and S15-S18. Each experiment group included at least three replicates and all the analysis was performed using IBM SPSS Statistics 25 and the Origin software was used for graph plotting.

## Supporting Information

Supporting Information is available from the Wiley Online Library or from the author.

## Acknowledgements

Z.F., S.B. and J.Q. contributed equally to this work. This work was supported by National Natural Science Foundation of China (61735016, 61975172, 61703365, 31871056, 91732302 and 21788102), Zhejiang Provincial Natural Science Foundation of China (LR17F050001), the National Key Research and Development Program of China (2018YFC1005003 and 2018YFE0190200), Fundamental Research Funds for the Central Universities (2020-KYY-511108-0007 and 2018QN81008), Research Grants Council of Hong Kong (C6009-17G and A-HKUST605/16) and Innovation and Technology Commission (ITC-CNERC14SC01 and ITCPD/17-9).

The procedures for all the animal experiments were approved by Animal Use and Care Committee at Zhejiang University (ZJU20190076) and in accordance with the National Institutes of Health Guidelines. All the experimenters have passed the medical laboratory animal training technology carried out by the Animal Center of Zhejiang University and obtained the qualified certificate.

We thank Dandan Song in the Center of Cryo-Electron Microscopy (CCEM), Zhejiang University for her technical assistance on Scanning Transmission Electron Microscopy.

## Notes

### Competing Interest Statement

The authors have declared no competing interest.

