## supplementary information for "Biologically excretable AIE dots for visualizing through the marmosets intravitally: horizons in future clinical nanomedicine"

Xiaoming Yu
Women's Hospital, Zhejiang University School of Medicine, Hangzhou 310006, China

Di Wu, Dr. Xiaoxiao Fan, Dr. Dingwei Xue
Sir Run Run Shaw Hospital, Zhejiang University School of Medicine, Hangzhou 310016, China

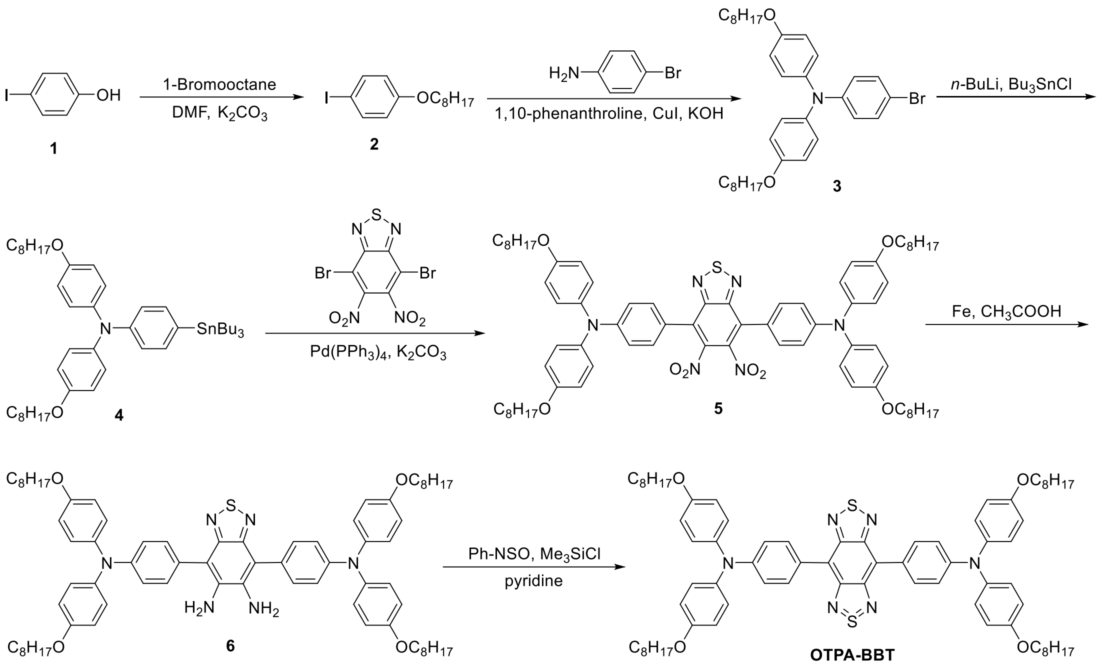


**Scheme S1.** Synthetic route to OTPA-BBT.

### Synthesis Processes and Characterizations

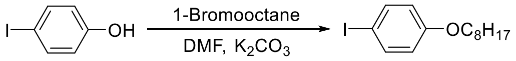


#### Synthesis of 1-iodo-4-(octyloxy)benzene (2)

1-Bromooctane (4.25 g, 22 mmol), 4-iodophenol (4.4 g, 20 mmol), and $K_2CO_3$ (8.28 g, 60 mmol) were added into a 250 mL of two-necked round-bottom flask, which was vacuumed and purged with dry nitrogen three times. Then anhydrous DMF (120 mL) was added, and the mixture was heated to reflux and stirred for 24 h. After cooling down to room temperature, water was added, and the mixture was washed with dichloromethane three times. The organic phase was combined, dried with $MgSO_4$, and the solvent was evaporated under reduced pressure. The crude product was purified by column chromatography on silica gel using dichloromethane/hexane (v/v 1:10) as the eluent to afford 1-iodo-4-(octyloxy)benzene as a colorless oil (83% yield). $^1$H NMR (400 MHz, $CDCl_3$): δ (ppm) 7.54 (d, 2H), 6.67 (d, 2H), 3.91 (t, 2H), 1.81-1.71 (m, 2H), 1.49-1.39 (m, 2H), 1.36-1.23 (m, 8H), 0.88 (t, 3H). $^{13}$C NMR (100 MHz, $CDCl_3$): δ (ppm) 159.03, 138.15, 116.94, 82.39, 68.14, 31.82, 29.34, 29.24, 29.16, 26.01, 22.67, 14.12.

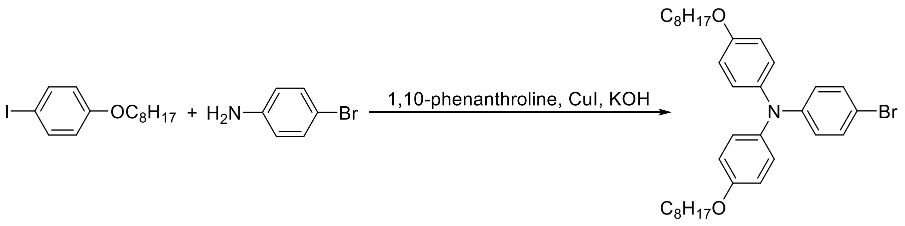


**Synthesis of 4-bromo-*N*,*N*-bis(4-(octyloxy)phenyl)aniline (3)**

4-Bromoaniline (1.03 g, 6 mmol), 1-iodo-4-(octyloxy)benzene (4.98 g, 15 mmol), 1,10-phenanthroline (0.18 g, 1 mmol), CuI (93 mg, 0.19 mmol), and KOH (5.04 g, 90 mmol) were added into a 250 mL of two-necked round-bottom flask. Then the flask was vacuumed and purged with dry nitrogen three times, and dry toluene (60 mL) was added into the flask, and the mixture was heated to reflux and stirred for 24 h. After cooling down to room temperature, water was added, and the mixture was washed with dichloromethane three times. The organic phase was combined, dried with $MgSO_4$, and the solvent was evaporated under reduced pressure. The crude product was purified by column chromatography on silica gel using dichloromethane/hexane (v/v 1:6) as the eluent to afford 4-bromo-*N*,*N*-bis(4-(octyloxy)phenyl)aniline as a viscous oil (76% yield). $^1$H NMR (400 MHz, $CDCl_3$): δ (ppm) 7.22 (d, 2H), 7.00 (d, 4H), 6.79 (t, 6H), 3.92 (t, 4H), 1.81-1.72 (m, 4H), 1.49-1.41 (m, 4H), 1.36-1.24 (m, 16H), 0.89 (t, 6H). $^{13}$C NMR (100 MHz, $CDCl_3$): δ (ppm) 155.65, 148.01, 140.36, 131.71, 126.57, 121.86, 115.33, 112.17, 68.28, 31.82, 29.37, 29.34, 29.25, 26.08, 22.66, 14.11.

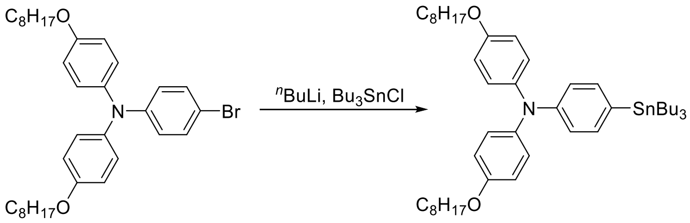


**Synthesis of 4-(octyloxy)-*N*-(4-(octyloxy)phenyl)-*N*-(4-(tributylstannyl)phenyl)aniline** (**4**)

4-Bromo-*N*,*N*-bis(4-(octyloxy)phenyl)aniline (2.32 g, 4 mmol) was added into a 100 mL of two-necked round-bottom flask. The flask was then vacuumed and purged with dry nitrogen three times, and anhydrous THF (45 mL) was added. Then the mixture was cooled with dry ice-acetone mixture to -78 $^o$C, and maintained at this temperature for 15 min, followed by the addition of *n*-butyllithium ($^n$BuLi, 2.5 M hexane solution, 1.6 mL, 4 mmol). After stirring at this temperature for 1.5 h, tri-*n*-butyltin chloride (1.2 mL, 4.5 mmol) was added, and the mixture was slowly warmed to room temperature, and stirred overnight. Afterwards, water was added to quench the reaction, and the mixture was extracted with dichloromethane three times. The organic phase was combined, dried with $MgSO_4$, and the solvent was evaporated under reduced pressure. The crude product was used without further purification.

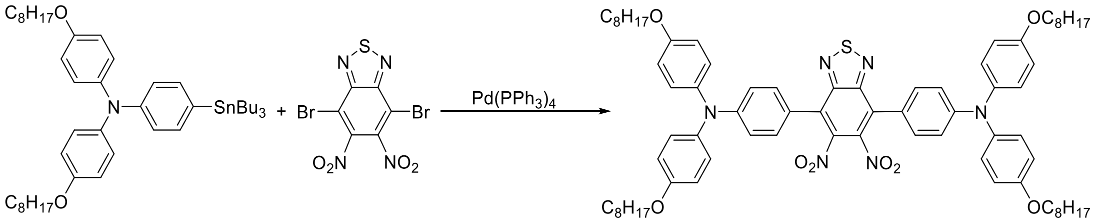


**Synthesis of 4,4'-(5,6-dinitrobenzo[*c*][1,2,5]thiadiazole-4,7-diyl)bis(*N*,*N*-bis(4-(octyloxy)phenyl)aniline) (5)**

4-(Octyloxy)-*N*-(4-(octyloxy)phenyl)-*N*-(4-(tributylstannyl)phenyl)aniline (2.37 g, 3 mmol), 4,7-dibromo-5,6-dinitrobenzo[*c*][1,2,5]thiadiazole (461 mg, 1.2 mmol), and $Pd(PPh_3)_4$ (70 mg, 0.06 mmol) were added into a 100 mL of two-necked round-bottom flask. The flask was then vacuumed and purged with dry nitrogen three times, and anhydrous THF (45 mL) was added. The mixture was heated to reflux and stirred for 24 h. After cooling down to room temperature, water was added, and the mixture was washed with dichloromethane three times. The organic phase was combined, dried with $MgSO_4$, and the solvent was evaporated under reduced pressure. The crude product was purified by column chromatography on silica gel using dichloromethane/hexane (v/v 1:2) as the eluent to afford 4,4'-(5,6-dinitrobenzo[*c*][1,2,5]thiadiazole-4,7-diyl)bis(*N*,*N*-bis(4-(octyloxy)phenyl)aniline) as a dark blue solid (71% yield). $^{1}$H NMR (400 MHz, $CDCl_3$): δ (ppm) 7.35 (d, 4H), 7.15 (d, 8H), 6.95 (d, 4H), 6.87 (d, 8H), 3.94 (t, 8H), 1.85-1.73 (m, 8H), 1.51-1.41 (m, 8H), 1.39-1.24 (m, 32H), 0.89 (t, 12H). $^{13}$C NMR (100 MHz, $CDCl_3$): δ (ppm) 156.49, 153.34, 150.63, 142.14, 139.25, 130.14, 127.91, 120.37, 117.73, 115.51, 68.28, 31.83, 29.38, 29.33, 29.26, 26.09, 22.68, 14.12. HRMS (MALDI-TOF, *m*/*z*) $[M]^+$ calcd for $C_{74}H_{92}O_8N_6S$, 1224.6697; found, 1224.6699.

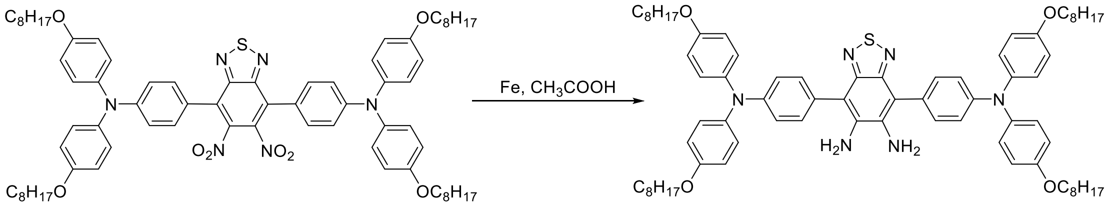


**Synthesis of 4,7-bis(4-(bis(4-(octyloxy)phenyl)amino)phenyl)benzo[*c*][1,2,5]thiadiazole-5,6-diamine (6)**

To the mixture of 4,4'-(5,6-dinitrobenzo[*c*][1,2,5]thiadiazole-4,7-diyl)bis(*N*,*N*-bis(4-(octyloxy)phenyl)aniline) (736 mg, 0.6 mmol) and acetic acid (80 mL) in a 250 mL of two-necked round-bottom flask, iron powder (1.01 g, 18 mmol) was added. The mixture was heated to 80 °C, and stirred for 3 h. After cooling down to room temperature, water was added, and the mixture was washed with dichloromethane three times. The organic phase was combined, dried with $MgSO_4$, and the solvent was evaporated under reduced pressure. The crude product was purified by column chromatography on silica gel using dichloromethane/hexane (v/v 2:1) as the eluent to afford 4,7-bis(4-(bis(4-(octyloxy)phenyl)amino)phenyl)benzo[*c*][1,2,5]thiadiazole-5,6-diamine as a dark red solid (75% yield). $^{1}$H NMR (400 MHz, $CDCl_3$): δ (ppm) 7.36 (d, 4H), 7.15 (d, 8H), 7.06 (d, 4H), 6.85 (d, 8H), 4.09 (s, 4H), 3.94 (t, 8H), 1.82-1.74 (m, 8H), 1.50-1.41 (m,

8H), 1.35-1.26 (m, 32H), 0.90 (t, 12H). $^{13}$C NMR (100 MHz, $CDCl_3$): δ (ppm) 155.78, 151.39, 148.51, 140.38, 138.14, 130.74, 127.21, 119.72, 115.34, 113.64, 68.28, 31.83, 29.38, 29.36, 29.25, 26.09, 22.67, 14.11.

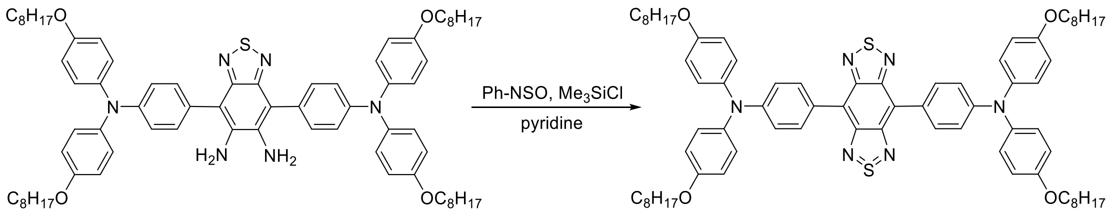


**Synthesis of 4,7-bis(4-(bis(4-(octyloxy)phenyl)amino)phenyl) benzo[1,2-*c*:4,5-*c*']bis[1,2,5]thiadiazole (OTPA-BBT)**

4,7-Bis(4-(bis(4-(octyloxy)phenyl)amino)phenyl)benzo[*c*][1,2,5]thiadiazole-5,6-diamine (466 mg, 0.4 mmol) was dissolved in dry pyridine (40 mL) in a 100 mL two-necked round-bottom flask. The flask was vacuumed and purged with dry nitrogen three times. Then *N*-sulfinylaniline (0.09 mL, 0.8 mmol) and trimethylsilyl chloride (0.15 mL, 1.2 mmol) were added into the solution. The mixture was heated to 80 °C, and stirred overnight. The solvent was evaporated under reduced pressure, and the residue was purified by column chromatography on silica gel using dichloromethane/hexane (v/v 1:2) as the eluent to result in 4,7-bis(4-(bis(4-(octyloxy)phenyl)amino)phenyl) benzo[1,2-*c*:4,5-*c*']bis[1,2,5]thiadiazole (OTPA-BBT) as a dark solid (66% yield). $^{1}$H NMR (400 MHz, $CDCl_3$, 25 °C): δ (ppm) 8.15 (d, 4H), 7.21 (d, 8h), 7.14 (d, 4H), 6.90 (d, 8H), 3.98 (t, 8H), 1.87-1.75 (m, 8H), 1.55-1.45 (m, 8H), 1.43-1.26 (m, 32H), 0.92 (t, 12H). $^{13}$C NMR (100 MHz, $CDCl_3$): δ (ppm) 156.07, 152.76, 149.27, 140.00, 132.59, 127.51, 126.64, 119.99, 118.62, 115.40, 68.31, 31.84, 29.39, 29.37, 29.26, 26.10, 22.67, 14.11. HRMS (MALDI-TOF, *m*/*z*) $[M]^+$ calcd for $C_{74}H_{92}O_4N_6S_2$, 1192.6621; found, 1192.6616.

**The measurement of the photoluminescence quantum yield.** A reference of IR-26 in 1,2-dichloroethane (DCE) with a nominal photoluminescence quantum yield of 0.5% was chosen to measure the NIR-II (beyond 900 nm) PLQY of OTPA-BBT dots in water. In order to reduce the attenuation of fluorescence in the material caused by the self-absorption of the material. Five dilute solutions with OD at 793 nm no more than 0.1 were prepared respectively. Their integrated emission intensities (900-1700 nm) were recorded under identical excitation conditions and the intensities were plotted as a function of OD at 793 nm for OTPA-BBT dots in DI water and IR-26 in DCE (reference solution). The quantum yield of OTPA-BBT dots was determined according to the comparison of the two slopes. The NIR-II quantum yield of the sample was calculated as follows:

$$PLQY = PLQY_{ref} \cdot \frac{F}{F_{ref}} \cdot \frac{A_{ref}}{A} \cdot \frac{I_{ref}}{I} \cdot \frac{n^2}{{n_{ref}}^2} = PLQY_{ref} \cdot \frac{Slope}{Slope_{ref}} \cdot \frac{n^2}{{n_{ref}}^2}$$

Where *PLQY* is the quantum yield of sample or reference, *F* is the integrated intensity, *A* is the absorption at 793 nm, *I* is the luminous intensity of the excitation source, *n* is the index of refraction of the solvent and *Slope* is the slope value in the Figure S9. Subscripts *ref* identify the reference. The Figure S2B-S2C showed the results of the photoluminescence quantum yield of the OTPA-BBT dots in water.

**Simulated intestinal fluid (SIF) preparation**: 6.8 g potassium dihydrogen phosphate was dissolved into 500 mL deionized water in a beaker. The pH of the solution was adjusted to 6.8 with sodium hydroxide solution (0.1 mol $L^{-1}$). 10 g pancreatin was dissolved into a moderate amount of deionized water in another beaker. The solutions were mixed and diluted with deionized water to 1000 mL for the formulation of SIF.

**Simulated gastric fluid (SGF) preparation**: 800 mL distilled water and 10 g pepsin were mixed into 16.4 mL diluted hydrochloric acid (HCl:$H_2O$, 234:766 vol/vol) and the mixture was stirred fully. The solution was then diluted to 1000 mL with deionized water for the formulation of SGF.

**Cerebrovascular microscopic imaging in mice**: 18-20 g male BALB/c mice were used for NIR-II fluorescence microscopic imaging of brain vasculatures. The mice were anesthetized with pentobarbital (50 mg $kgBW^{-1}$), and subsequently intravenously injected with AIE dotes (2 mg $kgBW^{-1}$). The images of cerebral vascular network in mice were recorded using NIR-II fluorescence microscopic imaging system. Before observation, a window of cerebral vessels was established by micro-craniotomy on each mouse, and a round thin coverslip was directly adhered to the mouse brain and secured by dental cement. The mouse's head was immobilized during the entire imaging process. The Figure S5 and S7 showed the results of NIR-II fluorescence microscopic cerebral vasculatures imaging in mice.

***In vivo* NIR-IIb GI tract imaging in mice**: OTPA-BBT dots (15 mg $kgBW^{-1}$) was perfused to the stomach of the nude mice (18-20 g). The treated mice were imaged with NIR-II fluorescence macroscopic imaging system.

**Toxicological analysis for mice**: Twelve mice were randomly divided into four groups (two control groups and two experiment groups). Each mouse in the experiment group was intravenously injected with OTPA-BBT dots (2 mg $kgBW^{-1}$) and each mouse in the control group was treated with sterile distilled water. At 1 d post treatment, blood samples were collected from mice in one control group (n = 3) and one experiment group (n = 3) for blood routine examination and blood biochemistry test. For each mouse, 0.3 mL blood sample was added into evacuated tubes with EDTA for blood routine examination, while 0.6 mL blood sample for blood biochemistry test was added into evacuated tubes without EDTA. After clotting, the samples for blood biochemistry test were separated by centrifugation (2000 rpm, 10 min). The serum was pipetted from the cellular elements and stored at 4 °C. Complete blood count (CBC) and blood biochemistry test were further conducted on a BC-2800 Vet Animal Auto Haematology Analyzer (MINDRAY), Chemray 240 and Chemray 800 Automated Biochemical Analyzers (Rayto Life and Analytical Sciences Co., Ltd., China.), respectively. At 28 d post treatment, we performed the same procedure described above on the other experimental group (n = 3) and the other control group (n = 3).

**Toxicological analysis for marmosets**: Blood samples (n = 3) were collected from three marmosets before injection. At 1st day and 4th week post injection of OTPA-BBT dots (2 mg kgBW$^{-1}$), the blood samples of them were collected again. The blood tests of the samples were same as those of mice.

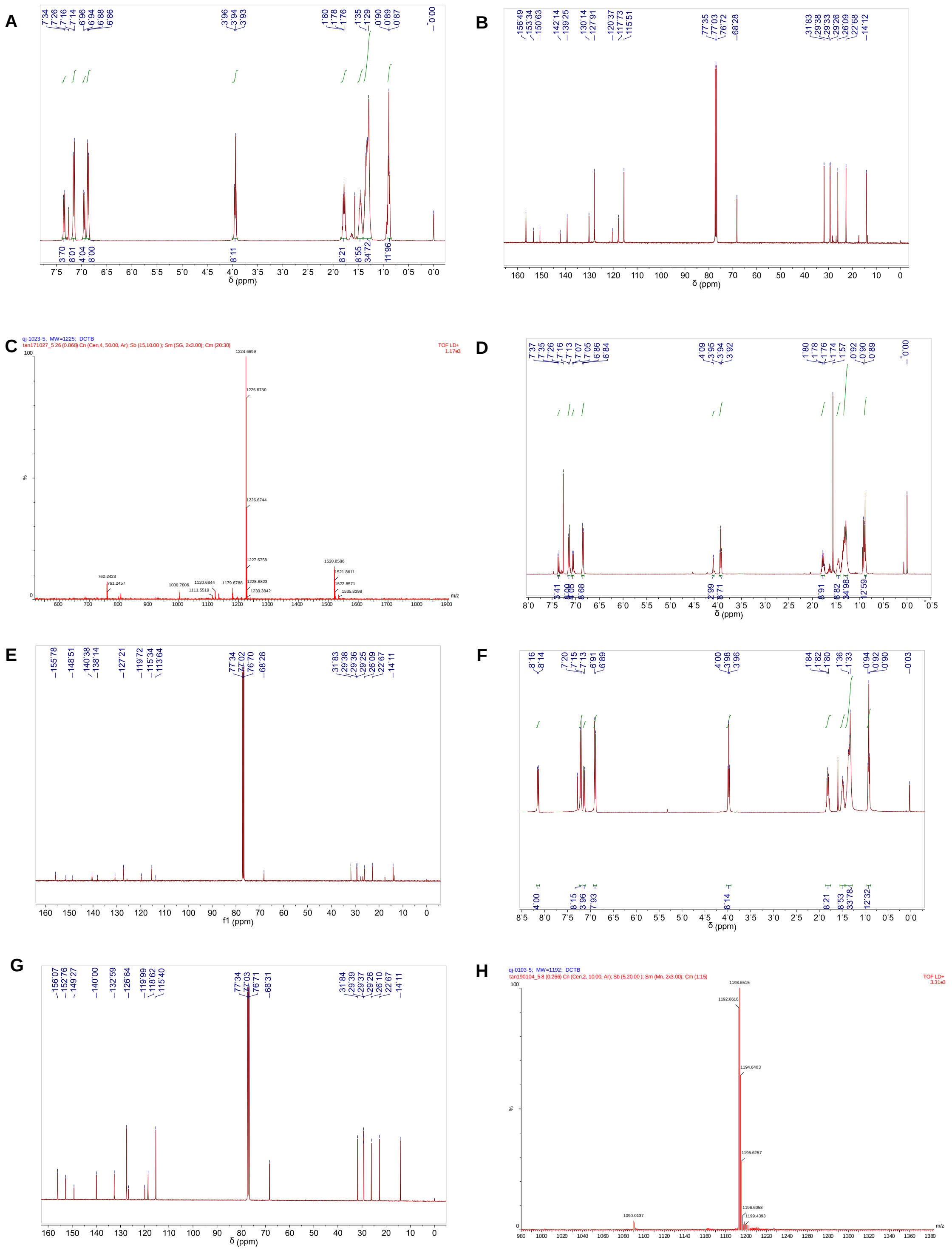

**Figure S1. Characterization of the products.** A) $^{1}$H NMR spectrum of 4,4'-(5,6-dinitrobenzo[*c*][1,2,5]thiadiazole-4,7-diyl)bis(*N*,*N*-bis(4-(octyloxy)phenyl)aniline) in $CDCl_3$ at 298 K. B) $^{13}$C NMR spectrum of 4,4'-(5,6-dinitrobenzo[*c*][1,2,5]thiadiazole-4,7-diyl)bis(*N*,*N*-bis(4-(octyloxy)phenyl)aniline) in $CDCl_3$ at 298 K. C) HRMS of 4,4'-(5,6-dinitrobenzo[*c*][1,2,5]thiadiazole-4,7-diyl)bis(*N*,*N*-bis(4-(octyloxy)phenyl)aniline). D) $^{1}$H NMR spectrum of 4,7-bis(4-(bis(4-(octyloxy)phenyl)amino)phenyl)benzo[*c*][1,2,5]thiadiazole-5,6-diamine in $CDCl_3$ at 298 K. E) $^{13}$C NMR spectrum of 4,7-bis(4-(bis(4-(octyloxy)phenyl)amino)phenyl)benzo[*c*][1,2,5]thiadiazole-5,6-diamine in $CDCl_3$ at 298 K. F) $^{1}$H NMR spectrum of OTPA-BBT in $CDCl_3$ at 298 K. G) $^{13}$C NMR spectrum of OTPA-BBT in $CDCl_3$ at 298 K. H) HRMS of OTPA-BBT.

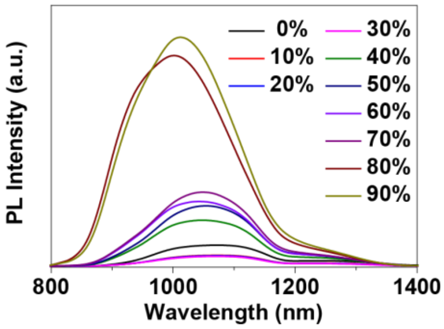


**Figure S2. PL spectra of OTPA-BBT in THF/$H_2O$ mixtures with different water fractions ($f_w$).**

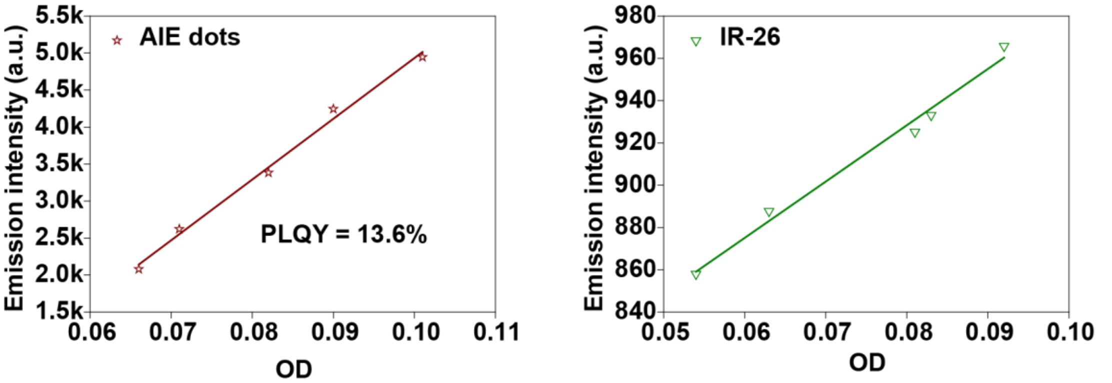


**Figure S3**. **The measurement of NIR-II photoluminescence quantum yield of OTPA-BBT dots, using IR-26 in dichloroethane as a reference.**

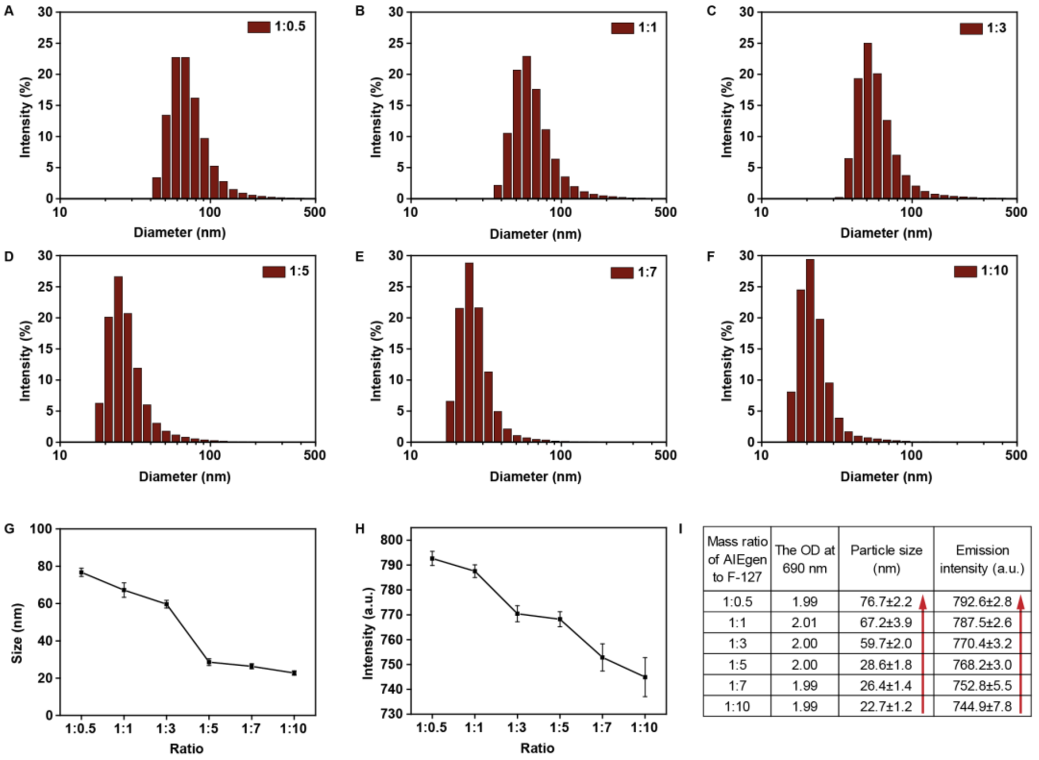


| Mass ratio of AIEgen to F-127 | The OD at 690 nm | Particle size (nm) | Emission intensity (a.u.) |
|---|---|---|---|
| 1:0.5 | 1.99 | 76.7±2.2 | 792.6±2.8 |
| 1:1 | 2.01 | 67.2±3.9 | 787.5±2.6 |
| 1:3 | 2.00 | 59.7±2.0 | 770.4±3.2 |
| 1:5 | 2.00 | 28.6±1.8 | 768.2±3.0 |
| 1:7 | 1.99 | 26.4±1.4 | 752.8±5.5 |
| 1:10 | 1.99 | 22.7±1.2 | 744.9±7.8 |

**Figure S4**. **The demonstration of AIE emission mechanism.** A) - F) The DLS results of OTPA-BBT dots dispersed in water with mass ratio of AIEgen and F-127 at A) 1:0.5, B) 1:1, C) 1:3, D) 1:5, E) 1:7 and F) 1:10. G) The mean particle sizes of OTPA-BBT dots with different mass ratios of AIEgen and F-127. H) The emission intensities of OTPA-BBT dots with different mass ratios of AIEgen and F-127 but the same OD at 690 nm wavelength (keeping the mass concentration of AIEgens the same). I) The table illustrated that fluorescence increased with the increase of particle size.

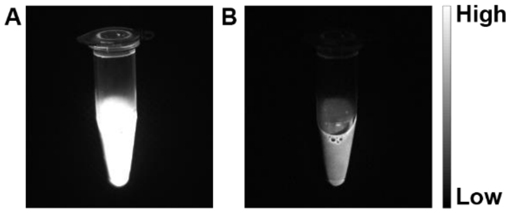


**Figure S5**. NIR-II fluorescence images of OTPA-BBT dots (0.8 mg $mL^{-1}$) beyond A) 1300 nm and B) 1500 nm under irradiation of 793 nm laser (50 mW $cm^{-2}$).

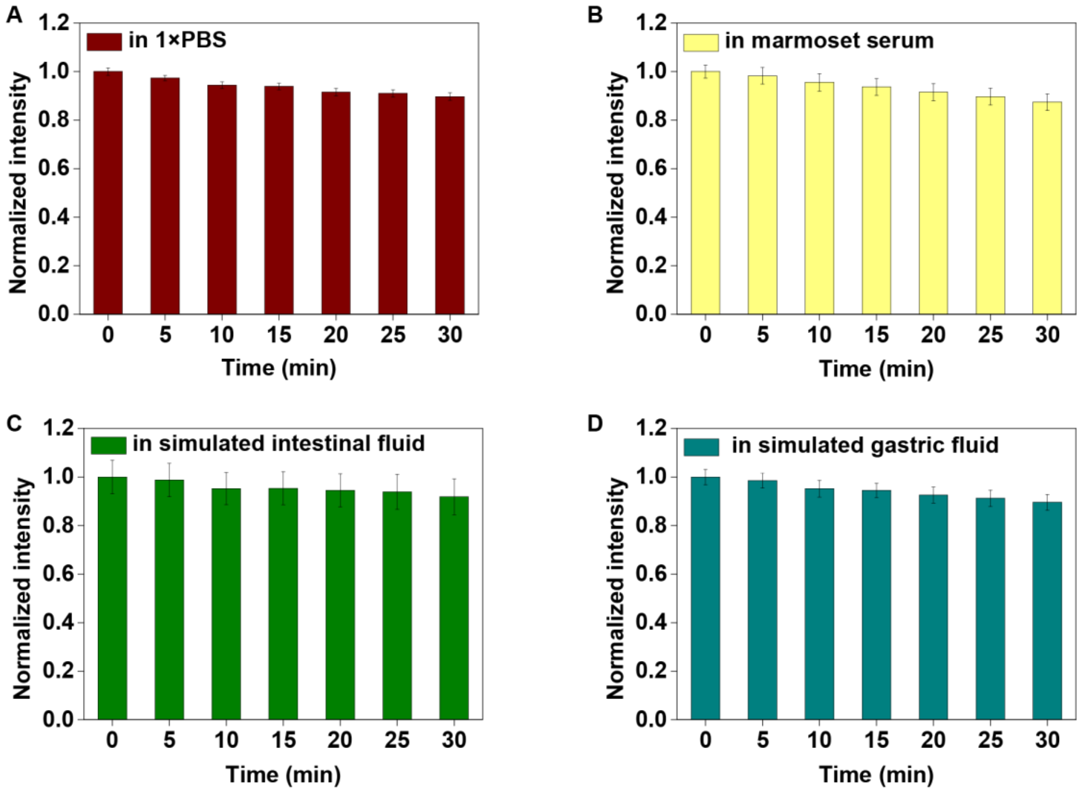


**Figure S6**. The photostability and chemical stability of the OTPA-BBT dots in A) phosphate buffer saline (PBS), B) marmoset serum, C) simulated intestinal fluid and D) gastric fluid, under the continuous irradiation of 793 nm CW laser (120 mW $cm^{-2}$) for 30 min.

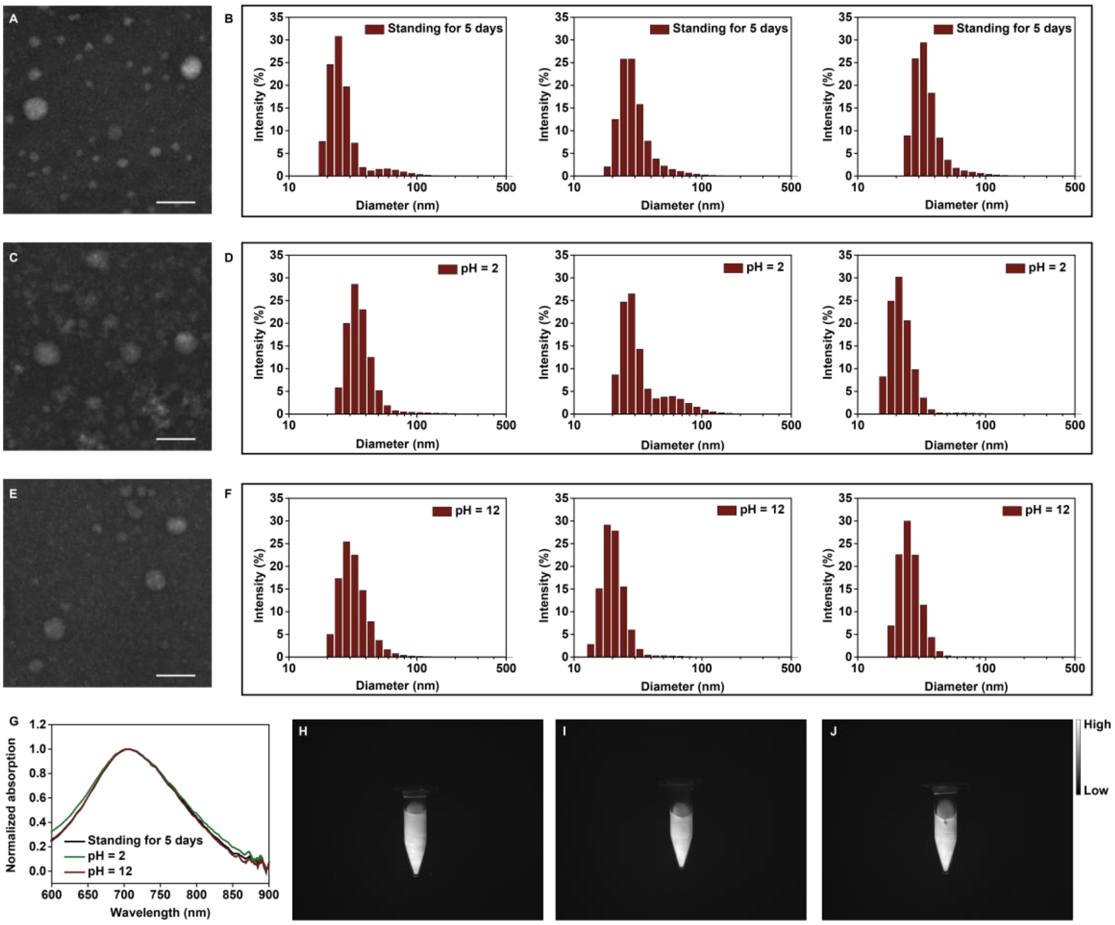


**Figure S7**. **The pH stability or long-term shelf-life stability of OTPA-BBT dots.** A) The STEM image and B) the DLS results (31.7 ± 2.3 nm) of OTPA-BBT dots that have been standing for 5 days. C) The STEM image and D) the DLS results (32.2 ± 4.7 nm) of OTPA-BBT dots in acidic environment (pH = 2) for 5 days. E) The STEM image and F) the DLS results (27.0 ± 3.6 nm) of OTPA-BBT dots in alkaline environment (pH = 12) for 5 days. G) The normalized absorption of OTPA-BBT dots that have been standing for 5 days, in acidic environment (pH = 2) for 5 days and in alkaline environment (pH = 12) for 5 days. The NIR-IIb fluorescence image of H) OTPA-BBT dots that have been standing for 5 days, I) in acidic environment (pH = 2) for 5 days and J) in alkaline environment (pH = 12) for 5 days taken under the same conditions. Scale bars in the STEM images represent 50 nm.

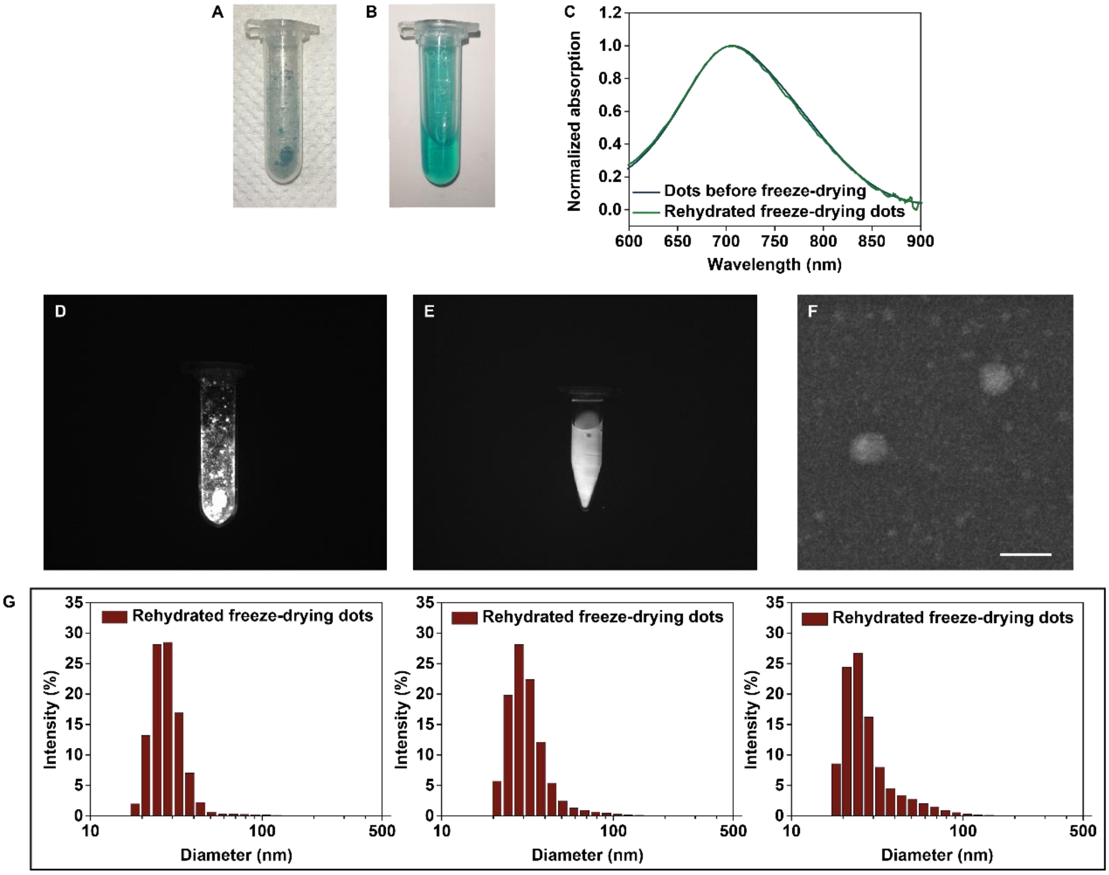


**Figure S8**. **The characterization of the freeze-dried and rehydrated dots.** (A) The picture of freeze-dried OTPA-BBT dots. (B) The picture of rehydrated OTPA-BBT dots after freeze-drying. (C) The normalized absorption spectra of OTPA-BBT dots before freeze-drying and the rehydrated OTPA-BBT dots. The NIR-IIb fluorescence images of (D) freeze-dried and (E) rehydrated OTPA-BBT dots. (F) The STEM image of the rehydrated dots. Scale bar, 50 nm. (G) The DLS results (30.3 ± 1.2 nm) of rehydrated OTPA-BBT dots.

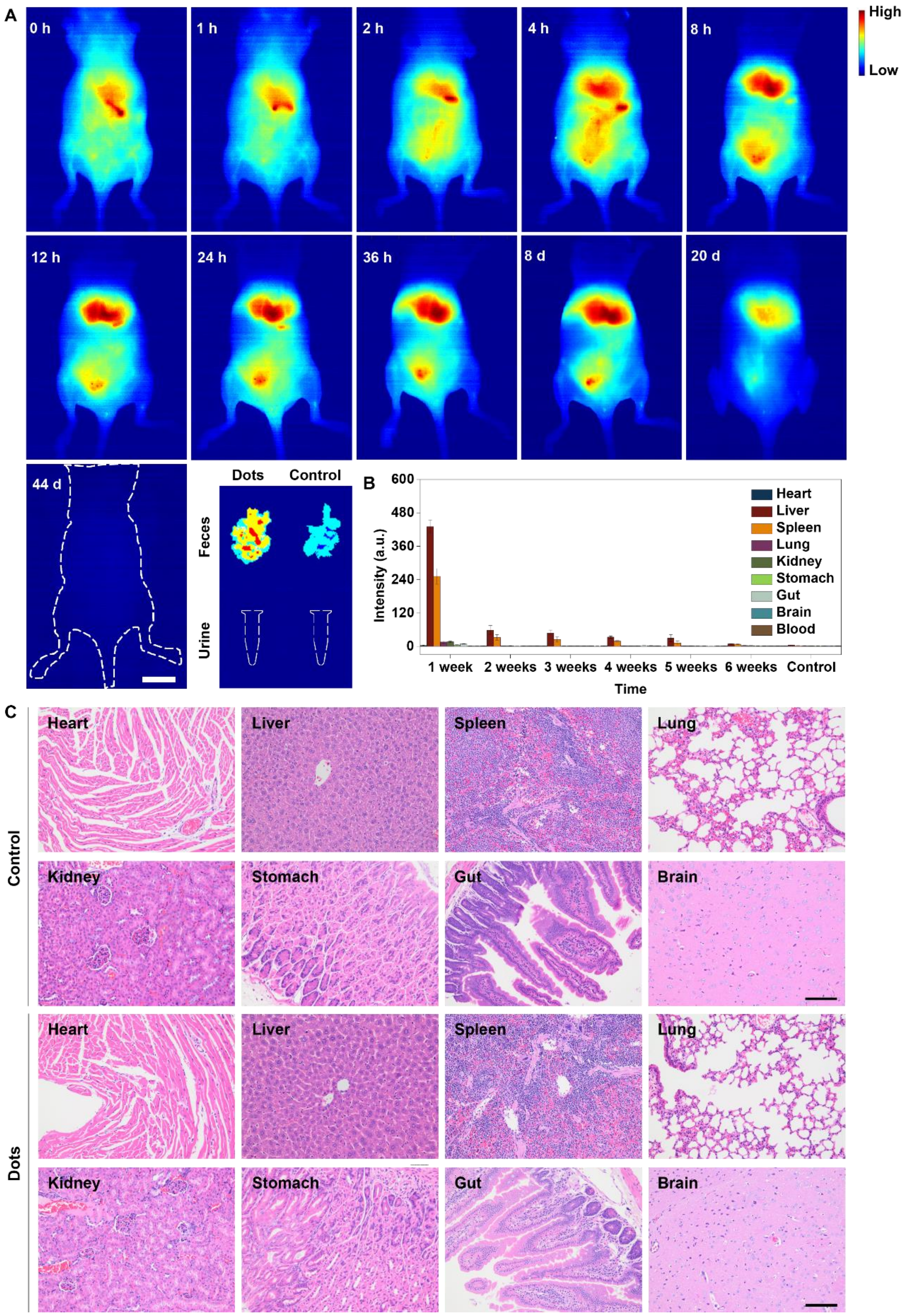


**Figure S9. The excretion and bio-toxicity of OTPA-BBT dots.** A) The *in vivo* excretion monitoring of OTPA-BBT dots in mice post intravenous injection (2 mg kgBW$^{-1}$), using the NIR-II fluorescence imaging modality. Scale bar, 10 mm. B) The biodistribution and excretion detection of OTPA-BBT dots *ex vivo* at various time post intravenous injection (2 mg kgBW$^{-1}$) on mice. C) Microscopic images of tissue sections from mice injected with 1 × PBS solution (200 μL) as control, and dispersion of OTPA-BBT dots (2 mg $kgBW^{-1}$) for 28 days. Scale bar: 100 μm.

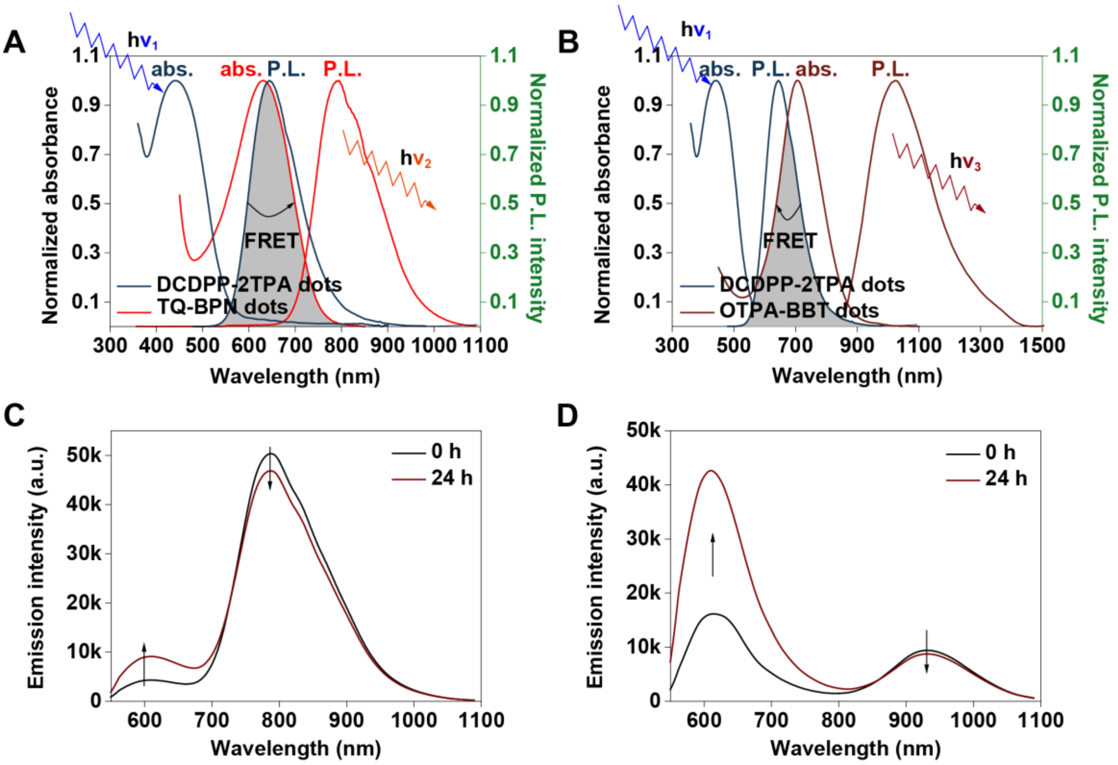


**Figure S10. The spectra of two kinds of FRET dots.** A) The absorption and emission spectra of DCDPP-2TPA dots and TQ-BPN dots. B) The absorption and emission spectra of DCDPP-2TPA dots and OTPA-BBT dots. The emission spectra of C) DCDPP2TPA-TQBPN dots and D) DCDPP2TPA-OTPABBT dots at 0 and 24 hours post mix with HSA.

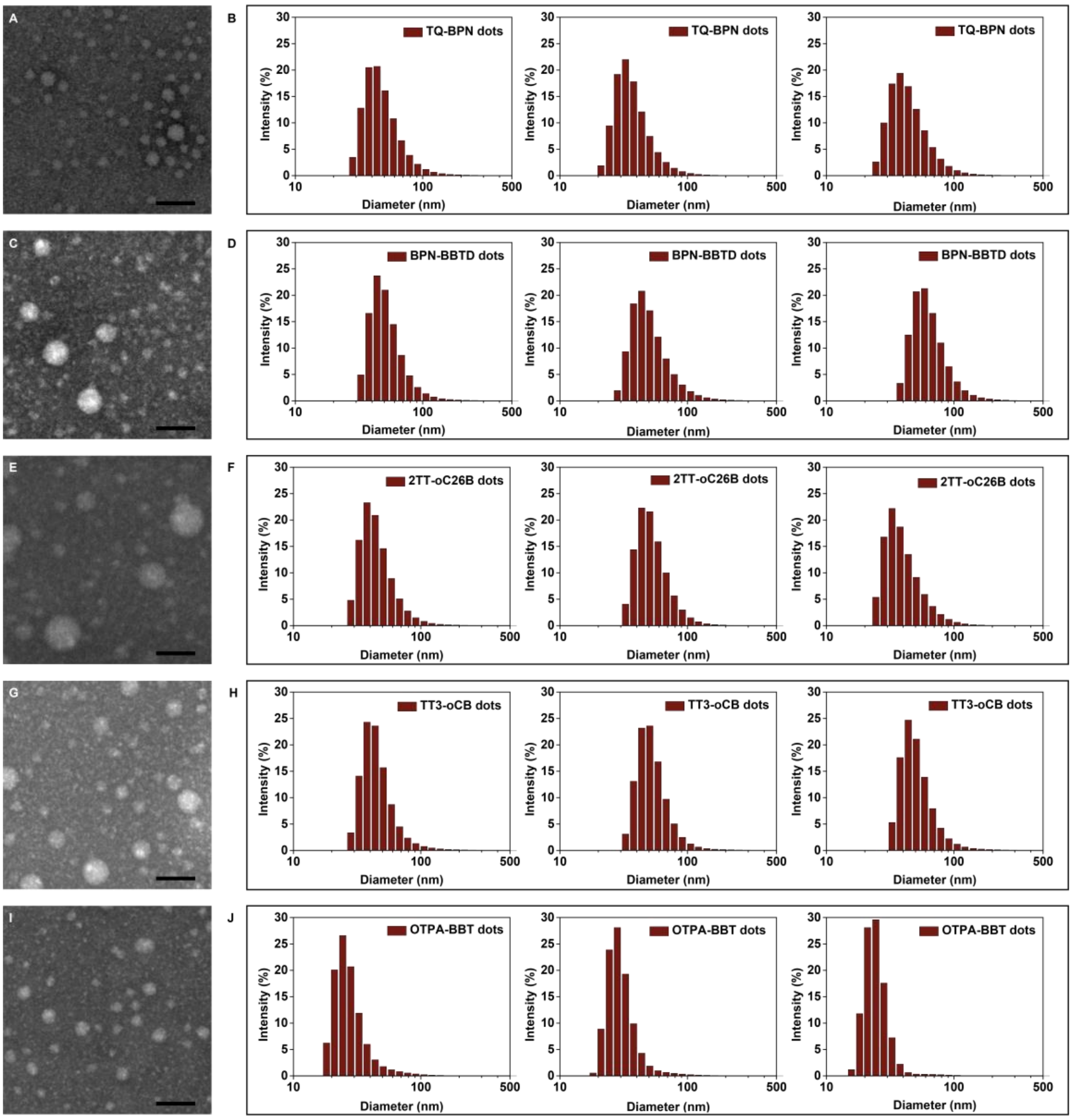


**Figure S11**. **The sizes and morphologies of several kinds of AIE dots.** A) The STEM image and B) the DLS results (44.9 ± 3.3 nm) of TQ-BPN dots. C) The STEM image and D) the DLS results (57.9 ± 4.2 nm). of BPN-BBTD dots E) The STEM image and F) the DLS results (47.6 ± 3.8 nm) of 2TT-oC26B dots. G) The STEM image and H) the DLS results (51.9 ± 2.4 nm) of TT3-oCB dots. I) The STEM image and J) the DLS results (28.6 ± 1.8 nm) of OTPA-BBT dots. Scale bars in the STEM images represent 50 nm.

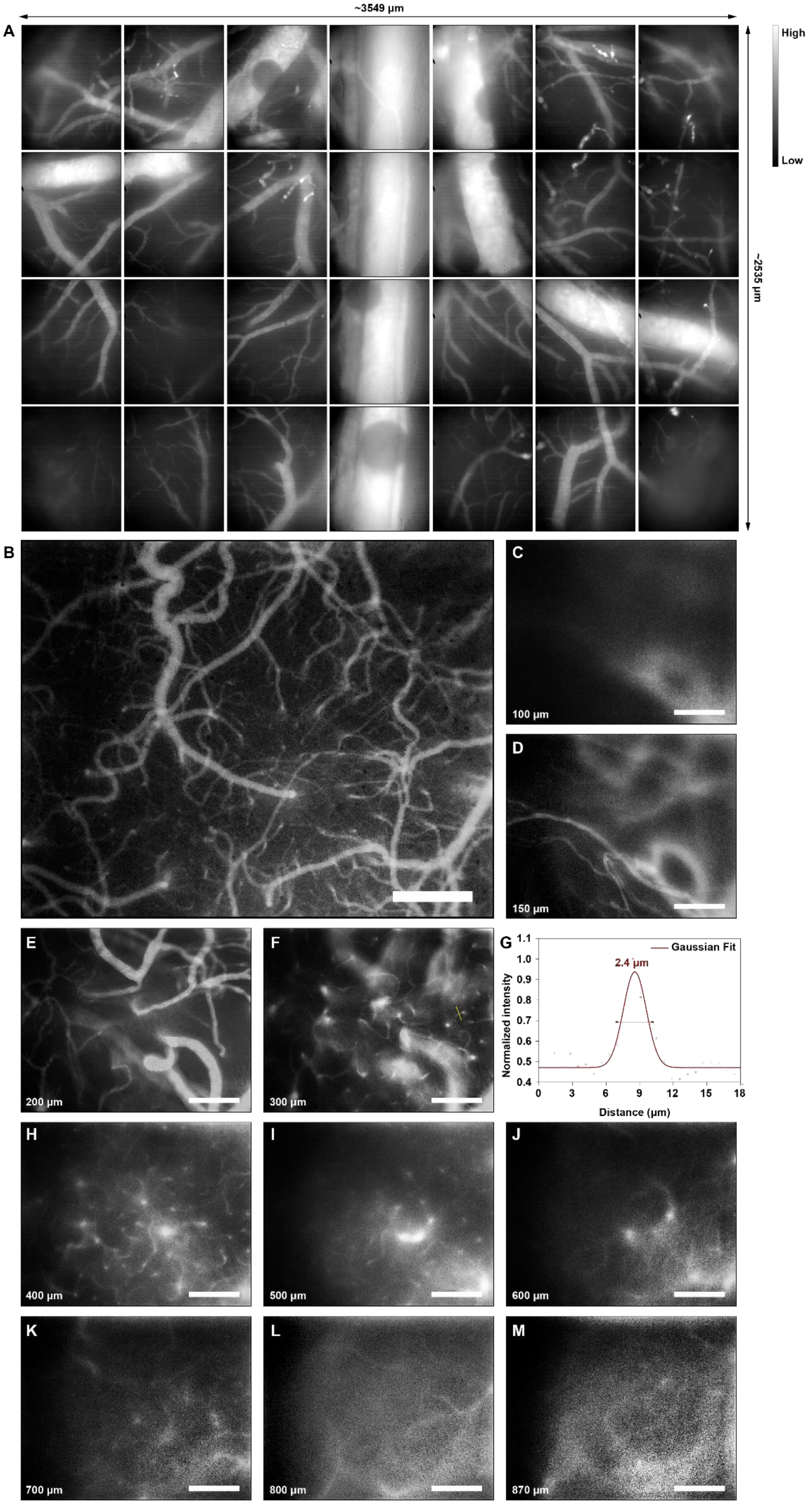
~3549 µm
A
High
Low
~2535 µm
B
C
100 µm
D
150 µm
E
200 µm
F
300 µm
G
Gaussian Fit
2.4 µm
Normalized intensity
Distance (µm)
H
400 µm
I
500 µm
J
600 µm
K
700 µm
L
800 µm
M
870 µm

**Figure S12. High-spatial-resolution NIR-II fluorescence microscopic cerebral vasculatures imaging in mice.** A) The NIR-II fluorescence cerebral vascular network under the cranial window of mice. The whole figure (with both large field-of-view and high spatial resolution) was stitched with 28 microscopic images (7×4) and each image was recorded by the 25X objective. B) A 5X NIR-II fluorescence microscopic image of vessels in mouse brain. Scale bar, 300 μm. C) - F) 25X images at various depths (100-300 μm) below the skull. Scale bar, 100 μm. G) Cross-sectional fluorescence intensity profile along the white line of the cerebral blood vessel (inset of (F)). The Gaussian fit to the profile is shown in red line in Figure S18G. H) - M) 25X images at various depths (400-870 μm) below the skull. Scale bar, 100 μm.

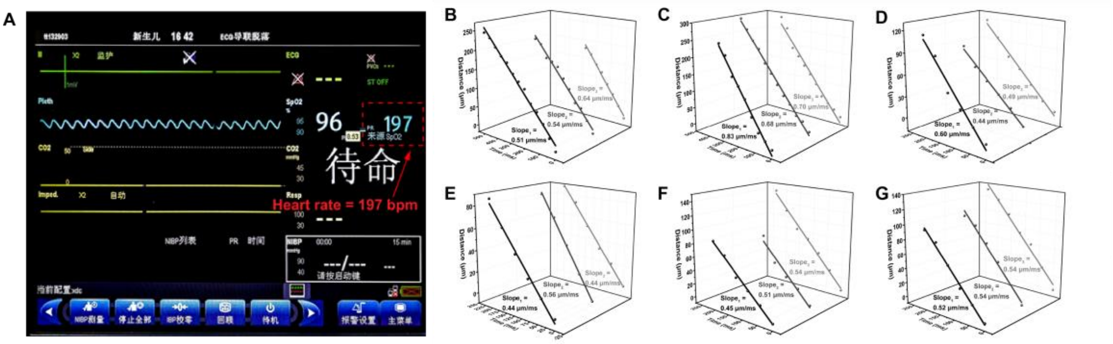


**Figure S13. The electrocardiogram and blood flow speeds of several brain vessels in marmoset.** A) The results of the ECG (electrocardiogram) monitor during cerebrovascular imaging in marmosets. The plots of positions of the point signal as a function of time (repeated 3 times) in the B) vessel 1, C) vessel 2, D) vessel 3, E) vessel 4, F) vessel 5 and G) vessel 6 in Figure 4A, respectively.

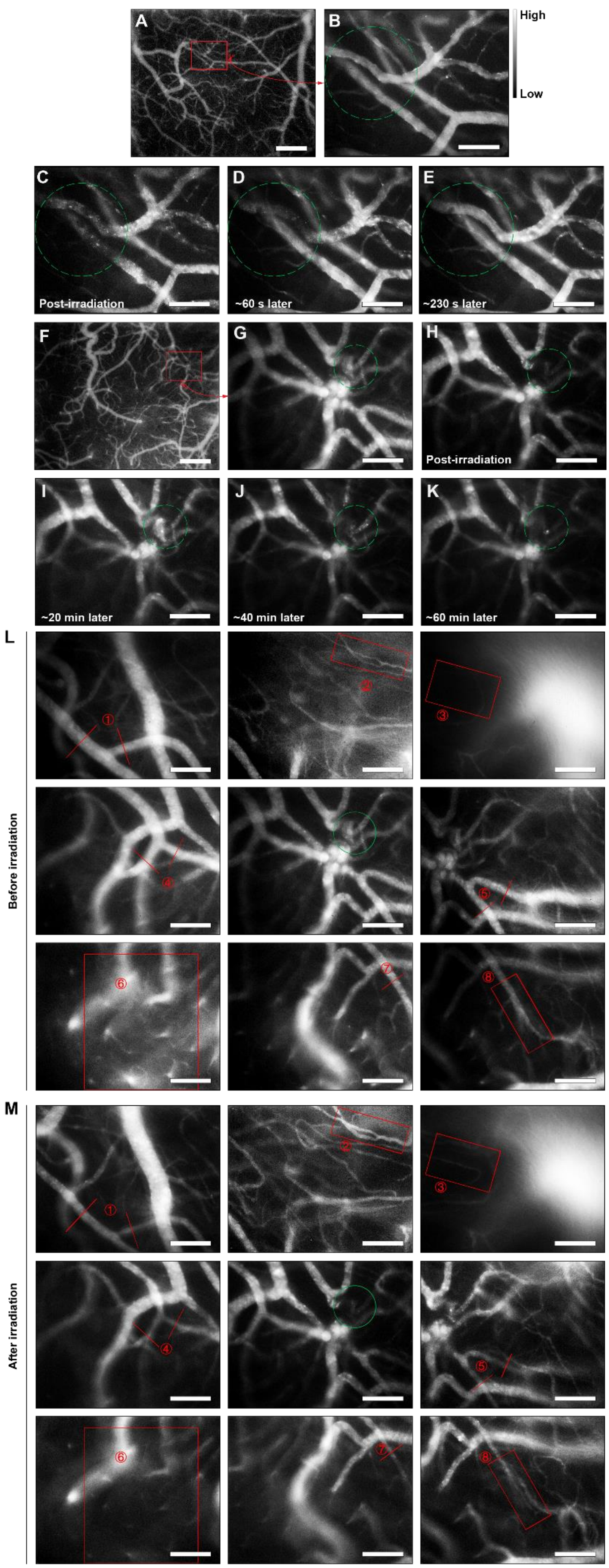

A
B
High
Low
C
D
E
Post-irradiation
~60 s later
~230 s later
F
G
H
Post-irradiation
I
J
K
~20 min later
~40 min later
~60 min later
L
Before irradiation
①
②
③
④
⑤
⑥
⑦
⑧
M
After irradiation
①
②
③
④
⑤
⑥
⑦
⑧

**Figure S14. The monitoring of the stokes in mice brain.** A) A large-field cerebrovascular image (5X) of a mouse. Scale bar, 300 μm. B) A selected region for further operation recorded by the 25X objective. The dashed green circle is the irradiation region of 532 nm CW laser. C) - E) The formation and recovery of the photothrombotic stroke during the 230 s post operation. Scale bar, 100 μm. F) A large-field cerebrovascular image (5X) of another mouse. Scale bar, 300 μm. G) A selected region for further operation recorded by the 25X objective. The dashed green circle is the irradiation region of 532 nm CW laser. H) - K) The formation of the photothrombotic stroke and the changes of the injured region during the 60 min post operation. Scale bar, 100 μm. The changes of the cerebral vasculatures L) before and M) after the photothrombotic stroke on blood vessels of nearby regions. (1), (4), (5), (6), (7) exhibited vasoconstriction. (2) exhibited vasodilation. (3), (8) exhibited hyperplasia. Scale bar, 100 μm.

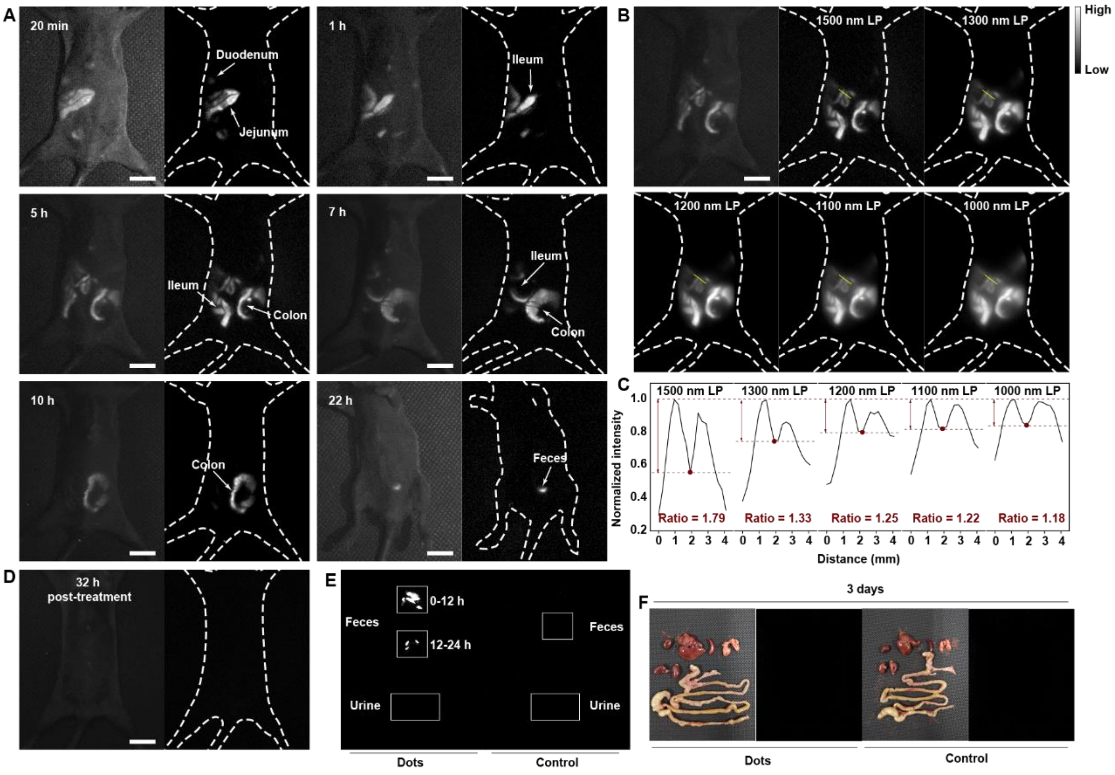


**Figure S15. NIR-IIb fluorescence GI imaging in mice.** A) NIR-IIb fluorescence GI imaging of mice, at various time post the intragastric administration of OTPA-BBT dots. Scale bar, 10 mm. B) Comparison of NIR-II fluorescence images beyond 1500 nm, 1300 nm, 1200 nm, 1100 nm, and 1000 nm. Scale bar, 10 mm. C) Cross-sectional fluorescence intensity profiles along the yellow lines (B) and the signal-to-background ratio of each profile where the background was defined as the valley between the two peaks. D) The image of the mouse at 32 h post treatment. Scale bar, 10 mm. E) The images of the feces and urine from the mice at various time post treatment. F) The image of the main organs of mice at 3 days post treatment.

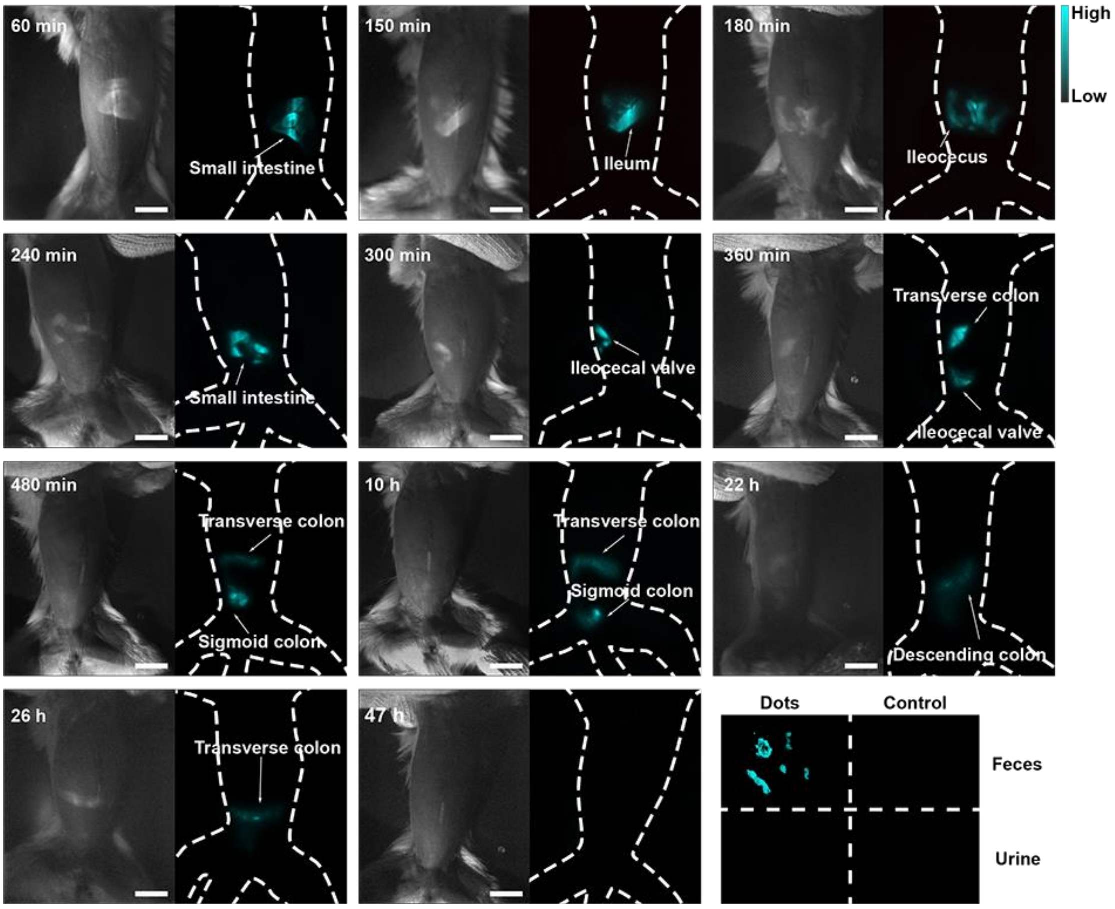


**Figure S16. The NIR-IIb fluorescence GI images and metabolite fluorescence test of a marmoset.** Scale bar, 20 mm.

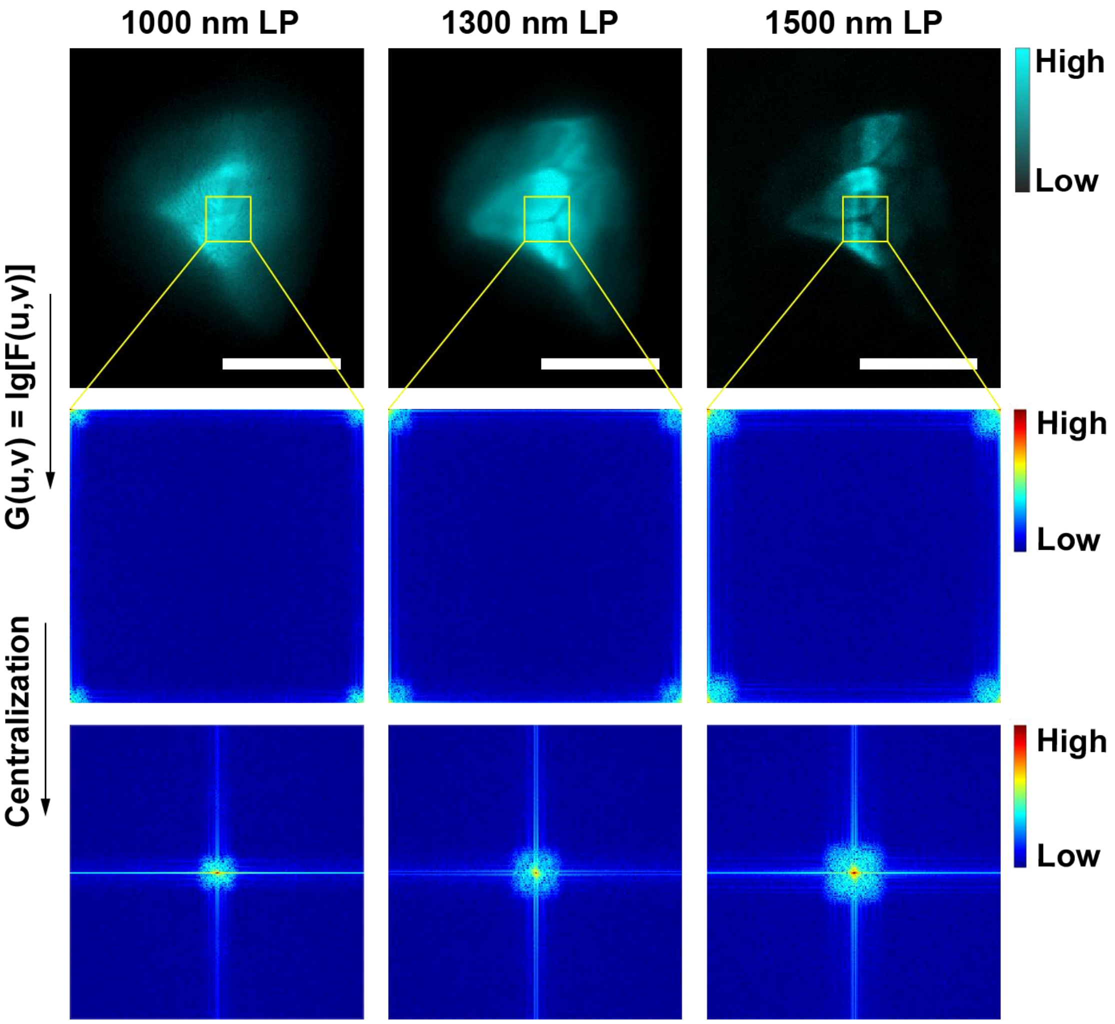


**Figure S17. The process of 2-D Fourier transform.** Scale bar, 20 mm.

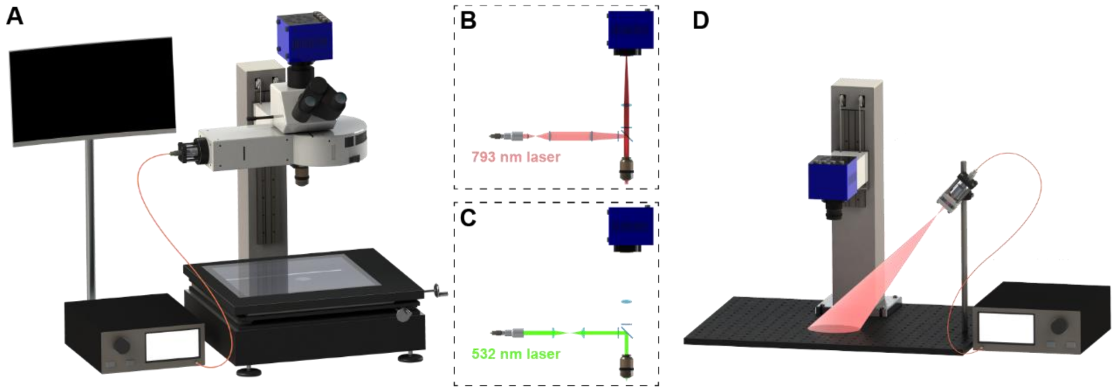


**Figure S18. Optical setup of the NIR-II fluorescence imaging.** A) Optical setup of the NIR-II fluorescence cerebrovascular microscopic monitoring. B) Model for imaging. C) Model for the induction of photothrombotic stroke. D) Optical setup of the NIR-II fluorescence GI imaging.

Tables S1-S8 showed the results of blood test for mice.

**Table S1. Results of blood routine examination for mice at 1st day post injection of sterile distilled water.**

| Index | Control sample 1 | Control sample 2 | Control sample 3 |
|---|---|---|---|
| White blood cell ($10^9$ $L^{-1}$) | 5.67 | 5.12 | 6.61 |
| Lymphocyte ($10^9$ $L^{-1}$) | 1.63 | 2.05 | 1.71 |
| Monocyte ($10^9$ $L^{-1}$) | 1.11 | 0.72 | 1.55 |
| Gran ($10^9$ $L^{-1}$) | 2.73 | 2.23 | 2.98 |
| Lymphocyte (%) | 28.7 | 40 | 25.8 |
| Monocyte (%) | 19.5 | 14 | 23.4 |
| Gran (%) | 48.3 | 43.7 | 45.2 |
| Red blood cell ($10^{12}$ $L^{-1}$) | 8.94 | 8.65 | 8.9 |
| Hemoglobin (g $L^{-1}$) | 155 | 152 | 156 |
| Hematocritg (%) | 37.7 | 36.3 | 37.6 |
| Mean corpuscular volume (fl) | 42.2 | 42 | 42.2 |
| Mean corpuscular hemoglobin (pg) | 17.3 | 17.6 | 17.5 |
| Mean corpuscular hemoglobin concentration (g $L^{-1}$) | 411 | 419 | 415 |
| Red cell volume Distribution Width (%) | 9.6 | 9.8 | 9.8 |
| Platelet ($10^9$ $L^{-1}$) | 358 | 449 | 507 |
| Mean platelet volume (fl) | 15.5 | 16.5 | 16.1 |
| Platelet distribution width | 16.7 | 16.4 | 16.5 |

**Table S2. Results of blood routine examination for mice at 1st day post injection of OTPA-BBT dots.**

| Index | Treated sample 1 | Treated sample 2 | Treated sample 3 |
|---|---|---|---|
| White blood cell ($10^9$ $L^{-1}$) | 5.28 | 6.45 | 5.27 |
| Lymphocyte ($10^9$ $L^{-1}$) | 1.81 | 1.9 | 1.89 |
| Monocyte ($10^9$ $L^{-1}$) | 0.8 | 1.01 | 0.84 |
| Gran ($10^9$ $L^{-1}$) | 2.62 | 3.43 | 2.48 |
| Lymphocyte (%) | 34.3 | 29.5 | 35.9 |
| Monocyte (%) | 15.2 | 15.7 | 15.9 |
| Gran (%) | 49.5 | 53.1 | 47.2 |
| Red blood cell ($10^{12}$ $L^{-1}$) | 8.93 | 8.86 | 8.26 |
| Hemoglobin (g $L^{-1}$) | 159 | 157 | 143 |
| Hematocritg (%) | 36.9 | 37.3 | 34 |
| Mean corpuscular volume (fl) | 41.3 | 42.1 | 41.2 |
| Mean corpuscular hemoglobin (pg) | 17.8 | 17.7 | 17.3 |
| Mean corpuscular hemoglobin concentration (g $L^{-1}$) | 431 | 421 | 421 |
| Red cell volume Distribution Width (%) | 9.6 | 9.6 | 9.9 |

| | | | |
|---|---|---|---|
| Platelet ($10^9$ $L^{-1}$) | 232 | 340 | 312 |
| Mean platelet volume (fl) | 17.1 | 15.2 | 15.3 |
| Platelet distribution width | 16.3 | 16.4 | 16.7 |

**Table S3. Results of hepatic and renal functions test for mice at 1st day post injection of sterile distilled water.**

| Index | Control sample 1 | Control sample 2 | Control sample 3 |
|---|---|---|---|
| ALT (U $L^{-1}$) | 65.3 | 60.2 | 67.6 |
| AST (U $L^{-1}$) | 135.4 | 136.2 | 134.2 |
| BUN (mmol $L^{-1}$) | 8.17 | 8.22 | 10.03 |
| Cre (μmol $L^{-1}$) | 36.4 | 30.6 | 33.7 |

**Table S4. Results of hepatic and renal functions test for mice at 1st day post injection of OTPA-BBT dots.**

| Index | Treated sample 1 | Treated sample 2 | Treated sample 3 |
|---|---|---|---|
| ALT (U $L^{-1}$) | 81.6 | 101.8 | 71 |
| AST (U $L^{-1}$) | 226.4 | 198.8 | 162.2 |
| BUN (mmol $L^{-1}$) | 7.94 | 7.82 | 8.24 |
| Cre (μmol $L^{-1}$) | 34.4 | 37.4 | 39.6 |

**Table S5. Results of blood routine examination for mice after 28 days post injection of sterile distilled water.**

| Index | Control sample 1 | Control sample 2 | Control sample 3 |
|---|---|---|---|
| White blood cell ($10^9$ $L^{-1}$) | 7.6 | 6.9 | 7.7 |
| Lymphocyte ($10^9$ $L^{-1}$) | 5.3 | 5.1 | 5.3 |
| Monocyte ($10^9$ $L^{-1}$) | 0.3 | 0.2 | 0.4 |
| Gran ($10^9$ $L^{-1}$) | 2 | 1.6 | 2 |
| Lymphocyte (%) | 69.5 | 73.1 | 69.4 |
| Monocyte (%) | 3.9 | 3.5 | 4.6 |
| Gran (%) | 26.6 | 23.4 | 26 |
| Red blood cell ($10^{12}$ $L^{-1}$) | 8.82 | 8.95 | 9.49 |
| Hemoglobin (g $L^{-1}$) | 114 | 117 | 125 |
| Hematocritg (%) | 37.4 | 37.5 | 39.9 |
| Mean corpuscular volume (fl) | 42.5 | 42 | 42.1 |
| Mean corpuscular hemoglobin (pg) | 12.9 | 13 | 13.1 |
| Mean corpuscular hemoglobin concentration (g $L^{-1}$) | 304 | 312 | 313 |
| Red cell volume Distribution Width (%) | 14.4 | 15.2 | 15.2 |
| Platelet ($10^9$ $L^{-1}$) | 999 | 873 | 912 |
| Mean platelet volume (fl) | 6.8 | 7.1 | 6.9 |
| Platelet distribution width | 16.1 | 16.6 | 16.3 |

**Table S6. Results of blood routine examination for mice after 28 days post injection of OTPA-BBT dots.**

| Index | Treated sample 1 | Treated sample 2 | Treated sample 3 |
|---|---|---|---|
| White blood cell ($10^9$ $L^{-1}$) | 7.6 | 7.4 | 6 |
| Lymphocyte ($10^9$ $L^{-1}$) | 5.5 | 5.2 | 3.8 |
| Monocyte ($10^9$ $L^{-1}$) | 0.3 | 0.3 | 0.2 |
| Gran ($10^9$ $L^{-1}$) | 1.8 | 1.9 | 2 |
| Lymphocyte (%) | 72.7 | 70.5 | 63.1 |
| Monocyte (%) | 3.5 | 3.9 | 3.9 |
| Gran (%) | 23.8 | 25.6 | 33 |
| Red blood cell ($10^{12}$ $L^{-1}$) | 9.23 | 8.82 | 9.75 |
| Hemoglobin (g $L^{-1}$) | 121 | 118 | 133 |
| Hematocritg (%) | 39.3 | 37 | 40.4 |
| Mean corpuscular volume (fl) | 42.6 | 42 | 41.5 |
| Mean corpuscular hemoglobin (pg) | 13.1 | 13.3 | 13.6 |
| Mean corpuscular hemoglobin concentration (g $L^{-1}$) | 307 | 318 | 329 |
| Red cell volume Distribution Width (%) | 15.5 | 15.2 | 15.9 |
| Platelet ($10^9$ $L^{-1}$) | 997 | 1020 | 574 |
| Mean platelet volume (fl) | 6.9 | 6.7 | 6.9 |
| Platelet distribution width | 16.3 | 16.4 | 16.4 |

**Table S7. Results of hepatic and renal functions test for mice after 28 days post injection of sterile distilled water.**

| Index | Control sample 1 | Control sample 2 | Control sample 3 |
|---|---|---|---|
| ALT (U $L^{-1}$) | 278.8 | 147.8 | 125.1 |
| AST (U $L^{-1}$) | 121.1 | 152.3 | 100.2 |
| BUN (mmol $L^{-1}$) | 4.71 | 4.09 | 4.45 |
| Cre (μmol $L^{-1}$) | 40.5 | 42.9 | 39.9 |

**Table S8. Results of hepatic and renal functions test for mice after 28 days post injection of OTPA-BBT dots.**

| Index | Treated sample 1 | Treated sample 2 | Treated sample 3 |
|---|---|---|---|
| ALT (U $L^{-1}$) | 146.6 | 200.1 | 236.2 |
| AST (U $L^{-1}$) | 118.7 | 126.7 | 116.3 |
| BUN (mmol $L^{-1}$) | 3.96 | 4.32 | 4.17 |
| Cre (μmol $L^{-1}$) | 40.8 | 38.1 | 44.1 |

Tables S9-S10 showed relative excretion statistics of five kinds of AIE dots.

**Table S9. Descriptive statistical results of five kind of AIE dots metabolisms.**

| Relative excretion ratio (RER)* | | 1 d | 2 d | 3 d | 4 d | 5 d | 6 d | 7 d |
|---|---|---|---|---|---|---|---|---|
| AIE dots without long aliphatic chains | TQ-BPN | 0.018 | 0.050 | 0 | 0.031 | 0.029 | 0.011 | 0.010 |
| | BPN-BBTD | 0.024 | 0.004 | 0 | 0.029 | 0.057 | 0.009 | 0.010 |
| AIE dots with long aliphatic chains | 2TT-oC26B | 0.018 | 0.147 | 0.052 | 0.017 | 0.037 | 0.034 | 0.020 |
| | TT3-oCB | 0.030 | 0.139 | 0.086 | 0.077 | 0.065 | 0.059 | 0.050 |
| | OTPA-BBT | 0.043 | 0.201 | 0.118 | 0.057 | 0.054 | 0.042 | 0.020 |

* Where relative excretion ratio (RER) $= \frac{I_{EX\text{-}feces} - I_{CTL\text{-}feces}}{I_{injected\ dots}}$. Once $I_{EX\text{-}feces} - I_{CTL\text{-}feces} < 0$, RER is set to zero.

**Table S10. Statistical difference between AIE dots with and without long aliphatic chains.**

| P value | AIE dots without long chains (n = 14) | AIE dots with long chains (n = 21) |
|---|---|---|
| AIE dots without long aliphatic chains | | 0.002 |
| AIE dots with long aliphatic chains | 0.002 | |

Tables S11-S14 showed the results of blood test for marmosets.

**Table S11. Results of blood routine examination for marmosets before the injection of OTPA-BBT dots.**

| Index | Before sample 1 | Before sample 2 | Before sample 3 |
|---|---|---|---|
| White blood cell ($10^9$ $L^{-1}$) | 10.95 | 6.72 | 4.48 |
| Lymphocyte ($10^9$ $L^{-1}$) | 2.29 | 1.17 | 0.97 |
| Monocyte ($10^9$ $L^{-1}$) | 1.87 | 1.11 | 0.95 |
| Gran ($10^9$ $L^{-1}$) | 6.44 | 4.28 | 2.6 |
| Lymphocyte (%) | 20.9 | 17.4 | 20.2 |
| Monocyte (%) | 17.1 | 16.5 | 20.2 |
| Gran (%) | 58.8 | 63.7 | 55.5 |
| Red blood cell ($10^{12}$ $L^{-1}$) | 7.02 | 7.56 | 5.85 |
| Hemoglobin (g $L^{-1}$) | 149 | 148 | 124 |

| | | | |
|---|---|---|---|
| Hematocritg (%) | 43.7 | 42.8 | 37.2 |
| Mean corpuscular volume (fl) | 62.2 | 56.6 | 63.4 |
| Mean corpuscular hemoglobin (pg) | 21.2 | 19.6 | 21.2 |
| Mean corpuscular hemoglobin concentration (g $L^{-1}$) | 341 | 346 | 333 |
| Red cell volume Distribution Width (%) | 8.2 | 9.3 | 9 |
| Platelet ($10^9$ $L^{-1}$) | 305 | 539 | 373 |
| Mean platelet volume (fl) | 12 | 11.1 | 12 |
| Platelet distribution width | 16.2 | 15.4 | 15.6 |

**Table S12. Results of blood routine examination for marmosets at 1st day post injection of OTPA-BBT dots.**

| Index | Before sample 1 | Before sample 2 | Before sample 3 |
|---|---|---|---|
| White blood cell ($10^9$ $L^{-1}$) | 9.95 | 6.26 | 6.14 |
| Lymphocyte ($10^9$ $L^{-1}$) | 1.45 | 0.73 | 1.78 |
| Monocyte ($10^9$ $L^{-1}$) | 1.78 | 1.34 | 0.9 |
| Gran ($10^9$ $L^{-1}$) | 6.39 | 4.05 | 3.36 |
| Lymphocyte (%) | 14.6 | 11.7 | 29 |
| Monocyte (%) | 17.9 | 21.4 | 14.6 |
| Gran (%) | 64.2 | 64.7 | 54.7 |
| Red blood cell ($10^{12}$ $L^{-1}$) | 7.44 | 5.95 | 7.02 |
| Hemoglobin (g $L^{-1}$) | 154 | 130 | 156 |
| Hematocritg (%) | 43.4 | 42.2 | 37 |
| Mean corpuscular volume (fl) | 61.8 | 56.7 | 62.2 |
| Mean corpuscular hemoglobin (pg) | 22.2 | 20.7 | 21.8 |
| Mean corpuscular hemoglobin concentration (g $L^{-1}$) | 359 | 365 | 351 |
| Red cell volume Distribution Width (%) | 9 | 9.7 | 9.1 |
| Platelet ($10^9$ $L^{-1}$) | 436 | 366 | 730 |
| Mean platelet volume (fl) | 12.7 | 11.2 | 11.6 |
| Platelet distribution width | 15.9 | 15.6 | 15.9 |

**Table S13. Results of hepatic and renal functions test for marmosets before the injection of OTPA-BBT dots.**

| Index | Before sample 1 | Before sample 2 | Before sample 3 |
|---|---|---|---|
| ALT (U $L^{-1}$) | 19.9 | 13.1 | 18.1 |
| AST (U $L^{-1}$) | 104.1 | 86.8 | 140.5 |
| BUN (mmol $L^{-1}$) | 4.09 | 6.07 | 4.31 |
| Cre (μmol $L^{-1}$) | 56 | 54 | 46.7 |

**Table S14. Results of hepatic and renal functions test for marmosets after 28 days post injection of OTPA-BBT dots.**

| Index | After sample 1 | After sample 2 | After sample 3 |
|---|---|---|---|
| ALT (U $L^{-1}$) | 11.9 | 12.5 | 9 |
| AST (U $L^{-1}$) | 78.8 | 89 | 99.1 |
| BUN (mmol $L^{-1}$) | 3.54 | 3.55 | 3.35 |
| Cre (μmol $L^{-1}$) | 41.5 | 53.2 | 39.6 |

Tables S15-S18 showed the statistical results of metabolites for marmosets.

**Table S15. Comparison of fluorescence intensity of marmoset feces after cerebrovascular imaging.**

| P value | Control (n = 5) | 1 d (n = 5) | 5 d (n = 5) | 10 d (n = 5) | 15 d (n = 5) | 26 d (n = 5) |
|---|---|---|---|---|---|---|
| Control | | 0 | 0 | 0 | 0.004 | 0.12 |
| 1 d | | | 0 | 0 | 0 | 0 |
| 5 d | | | | 0.161 | 0.021 | 0.001 |
| 10 d | | | | | 0.3 | 0.035 |
| 15 d | | | | | | 0.225 |
| 26 d | | | | | | |

**Table S16. Comparison of fluorescence intensity of marmoset urine after cerebrovascular imaging.**

| P value | Control (n = 5) | 1 d (n = 5) | 5 d (n = 5) | 10 d (n = 5) | 15 d (n = 5) | 26 d (n = 5) |
|---|---|---|---|---|---|---|
| Control | | 0.901 | 0.775 | 0.334 | 0.18 | 0.474 |
| 1 d | | | 0.682 | 0.286 | 0.222 | 0.553 |
| 5 d | | | | 0.505 | 0.108 | 0.32 |
| 10 d | | | | | 0.028 | 0.104 |
| 15 d | | | | | | 0.52 |
| 26 d | | | | | | |

**Table S17. Comparison of fluorescence intensity of marmoset feces after gastrointestinal imaging.**

| P value | Control (n = 3) | 0-6 h (n = 3) | 6-24 h (n = 3) | 24-42 h (n = 3) | 42-60 h (n = 3) |
|---|---|---|---|---|---|
| Control | | 0 | 0 | 0 | 0.252 |
| 0-6 h | | | 0.415 | 0 | 0 |
| 6-24 h | | | | 0 | 0 |
| 24-42 h | | | | | 0 |
| 42-60 h | | | | | |

**Table S18. Comparison of fluorescence intensity of marmoset urine after gastrointestinal imaging.**

| P value | Control (n = 3) | 0-1 d (n = 3) | 1-1.5 d (n = 3) | 1.5-2 d (n = 3) |
|---|---|---|---|---|
| Control | | 0.666 | 0.794 | 0.326 |
| 0-1 d | | | 0.863 | 0.173 |
| 1-1.5 d | | | | 0.224 |
| 1.5-2 d | | | | |

**Movie S1. The video of periodic vibrations of the capillaries.** Scale bar, 100 μm.

**Movie S2. The video of blood flow in marmosets.** Scale bar, 100 μm.

**Movie S3. The recovery of the photothrombotic micro-stroke of one vessel in mouse brain.** Scale bar, 100 μm.

**Movie S4. The recovery of the photothrombotic micro-stroke of the other vessel in mouse brain.** Scale bar, 100 μm.

**Movie S5. The record of the blood flow arrest or even reflux in marmosets.** Scale bar, 100 μm.

**Movie S6. The video of gastrointestinal peristalsis in mice at ~7 h post treatment.** Scale bar, 10 mm.

**Movie S7. The video of gastrointestinal peristalsis in marmosets at ~30 min post treatment.** Scale bar, 20 mm.
